# Connectomic and behavioral alterations in creatine transporter deficiency are partially normalized by gene therapy

**DOI:** 10.1101/2024.01.12.575377

**Authors:** Caterina Montani, Ludovica Iovino, Federica Di Vetta, Jean-Charles Rene’ Pasquin Mariani, A. Elizabeth De Guzman, Silvia Gini, Alberto Galbusera, Bianca D’Epifanio, Elsa Ghirardini, Sara Cornuti, Lorenzo Dadà, Elena Putignano, Maria Grazia Alessandrì, Giulia Vasirani, Sine Mandrup Bertozzi, Andrea Armirotti, Laura Baroncelli, Alessandro Gozzi

**Affiliations:** Functional Neuroimaging Laboratory, Istituto Italiano di Tecnologia, Center for Neuroscience and Cognitive Systems, CNCS@UNITN, Rovereto, Italy; Institute of Neuroscience, National Research Council (CNR), Pisa, Italy; Department of Biology, University of Pisa, Pisa, Italy; Department of Developmental Neuroscience, IRCCS Stella Maris Foundation, Pisa, Italy; BIO@SNS lab, Scuola Normale Superiore di Pisa, Pisa, Italy; Department of Statistics, Computer Science, Applications “Giuseppe Parenti” (DiSIA), University of Florence, Florence, Italy; University of Genova, Genova, Italy; IRCCS Ospedale Policlinico San Martino, Genova, Italy; Analytical Chemistry Facility, Istituto Italiano di Tecnologia, Genova, Italy

**Author notes:** Correspondence to: Alessandro Gozzi Functional Neuroimaging Laboratory, Istituto Italiano di Tecnologia, Center for Neuroscience and Cognitive Systems, CNCS@UNITN, Rovereto, Italy, Caterina Montani IRCCS Ospedale Policlinico San Martino, Genova, Italy. Current address: IRCCS Ospedale Policlinico San Martino, Genova, Italy (Caterina Montani). Contributed equally (Caterina Montani, Ludovica Iovino, Federica Di Vetta). Contributed equally (Laura Baroncelli, Alessandro Gozzi).

## Abstract

Creatine Transporter Deficiency (CTD) is an X-linked disorder due to the loss of *SLC6A8* gene and presenting with low brain creatine, intellectual disability, autistic-like behavior and seizures. No treatments are available yet for CTD, and little is known about the brain circuit alterations underlying its pathological endophenotypes. Here, we tracked brain network and behavioral dysfunction in a murine model of CTD at two stages of disease progression. fMRI mapping revealed widespread disruption of brain connectivity in Slc6a8-KO mice, with robust somatomotor hypoconnectivity in juvenile animals, and weaker and more focal alterations of cortical and subcortical connectivity in adulthood. Notably, perinatal AAV-mediated expression of human SLC6A8 in Slc6a8- KO mice robustly prevented juvenile fMRI hypoconnectivity, an effect accompanied by the regression of multiple translationally relevant phenotypes, including reduced stereotyped movements, improved declarative memory and increased body weight, all of which persisted into adulthood. However, early cognitive deficits, impairments in working memory and residual fMRI hypoconnectivity in adult mice were not ameliorated by gene therapy. Furthermore, significant cognitive impairments were observed in WT mice receiving gene therapy, highlighting a potential detrimental effect of ectopic expression of SLC6A8 in healthy brain circuits. Finally, multivariate modeling in adult mice revealed a basal forebrain network whose activity was associated with behavioral performance and modulated by brain creatine levels. This brain-behavior relationship was disrupted in Slc6a8-KO mice. Our results document robust network disruption in CTD and demonstrate that CTD pathology can be partially alleviated by perinatal genetic expression of SLC6A8, providing a foundation for the future development of experimental therapies for this genetic disorder.

## Introduction

Creatine Transporter Deficiency (CTD, OMIM: #300352) is an X-linked disease originating from mutations of the solute carrier family 6-member 8 (*SLC6A8*) gene^1^. *SLC6A8* encodes the protein responsible for cellular creatine (Cr) uptake. The clinical manifestation of CTD follows lack of Cr in the brain and is predominantly neurological. It includes global developmental delay, mild to severe intellectual disability (ID), disturbance of expressive and cognitive speech, psycho-motor impairment, autistic-like behavior, and seizures^2,3^. Despite our increased understanding of the etiopathological cascade underlying CTD^3,4^, effective treatments for this monogenic disorder are lacking. Dietary supplementation with Cr, either alone or in combination with its synthesis precursors, has shown very limited success^2,3^. Indeed, loss-of-function of *SLC6A8* not only prevents transport of Cr across the plasmatic membrane, but also affects its endogenous synthesis, which requires the coordinated action of multiple cell populations^5,6^. As a result, the current standard of care for CTD only involves pharmacological control of epilepsy and other CTD symptomatology.

Over the last few years, studies in murine models have revealed that neurological damage produced by CTD does not stem from overt neuronal death or degeneration, but it rather reflects more subtle alterations in synaptic compartments. Specifically, our own work previously documented a loss of GABAergic synapses in the cerebral cortex of mice lacking *Slc6a8* (Slc6a8-KO). This effect was associated with a dysfunction of parvalbumin-expressing GABAergic interneurons^7,8^. Supporting a key etiopathological role of altered inhibitory transmission in CTD, we also described marked abnormalities in EEG neural oscillations, with decreased theta/alpha power and increased gamma activity in both Slc6a8-KO mice and CTD patients^9^. Owing to the key contribution of oscillatory activity in coordinating large-scale patterns of brain synchronization and interareal communication^10^, this observation suggests that the neurological alterations that characterize CTD may be partly due to impaired long-range functional connectivity^11^. This notion is consistent with neuroimaging observations in CTD patients, where abnormalities in anatomical connectivity (including white matter demyelination, and corpus callosum thinning) have been reported^12^.

Here, we used fMRI connectivity mapping^13,14^ and behavioral testing in Slc6a8- KO mice to (i) longitudinally track the neural circuits primarily affected in CTD and (ii) relate connectivity alterations to the behavioral deficits that characterize this disorder. Importantly, we also probed whether perinatal adeno-associated virus (AAV)-mediated expression of a functional *SLC6A8* transgene could restore proper creatine transporter (CRT) expression and Cr level in the brain, thereby averting the manifestations of connectomic and behavioral phenotypes. Our results show that CTD is associated with distinct alterations in functional connectivity that evolve along its pathological trajectory. We also found that early intracerebral expression of human *SLC6A8* (hSLC6A8) may partly ameliorate both fMRI hypoconnectivity and behavioral impairment in Slc6a8-KO mice. These results provide proof-of-concept evidence that CTD phenotypes may be partly reversed by genetic therapies aimed at reinstating homeostatic Cr levels.

## Materials and Methods

Full-length experimental procedures can be found in the online Supplementary material.

### Ethical statement

Animal research was conducted in agreement with the Italian Law (DL 26/2014 of Italian Ministry of Health implementing EU 63/2010) and the National Institute of Health. Animal projects were approved by the Animal Care Committee of the University of Trento, Istituto Italiano di Tecnologia and Neuroscience Institute of Consiglio Nazionale delle Ricerche.

### Animals

Slc6a8-KO mice were generated by the European Molecular Biology Laboratory (EMBL). Slc6a8^x/-^ females were crossed with wild-type male mice to generate Slc6a8^-/y^ males (Slc6a8-KO, hereafter KO). Slc6a8^+/y^ wild-type male littermates were used as controls (WT). Mice were maintained in a controlled environment with humidity set at 60±10% and temperature maintained at 21±1 °C. Food and water were readily accessible ad libitum. Each standard cage provided nesting material and accommodated from 2 to 5 mice.

### Molecular cloning

The full-length human sequence of *SLC6A8* (*hSLC6A8*) was cloned under the control of the small JeT promoter^16^ into a pAAV_WPRE.SV40. An HA tag at the C-terminal of *SLC6A8* was used to detect the transgenic protein (pAAV_JeT-hSLC6A8- HA_WPRE.SV40). The JeT-hSCL6A8 sequence was synthesized by Twin Helix (Milano, Italy) as reported in Table S1. To verify that the HA tag does not interfere with transporter function, we also generated a version of the pAAV_JeT- hSLC6A8_WPRE.SV40 plasmid without the HA tag.

### Cell culture and transfection

HEK293T cells were maintained in DMEM with 10% FBS and 1% penicillin/streptomycin. Cell cultures were incubated at 37°C in a humidified atmosphere with 5% CO . For analysis of creatine (Cr) levels, upon reaching ∼80% confluency, cells were transfected using Lipofectamine 2000 at a 1:2 DNA/Lipofectamine ratio. Each well received 1.5 µg of plasmid DNA (JeT-hSLC6A8 or JeT-hSLC6A8-HA). The following day, cells were stimulated for 24 h with 125µM Cr. After treatment, the cells were centrifuged and pellets were stored at -80°C.

### Viral preparation

Serotype 9 adeno-associated viral (AAV) vectors containing pAAV_JeT- hSLC6A8-HA_WPRE.SV40 were produced by the University of Pennsylvania Vector Core (Philadelphia, PA). As control, we used an AAV9 containing pAAV_JeT- GFP_WPRE.SV40.

### Experimental design

Investigations at IIT (Gozzi laboratory, Rovereto, Italy) were carried out on four groups of male mice as follows: Slc6a8-KO (n=20) and WT (n=10) injected with AAV- hSLC6A8, and Slc6a8-KO (*n*=15) and WT (*n*=23) injected with AAV-GFP as controls. The four groups of mice were administered with AAV vectors via intracerebroventricular (i.c.v.) injection at P1. The four experimental groups were subjected to fMRI scans, at P40 and P140. Forty-eight hours after each fMRI session mice underwent two behavioral tests. The testing order consisted of self-grooming scoring, followed by the Y maze test. The rest of the experimental mice were used for post-mortem Cr quantifications, using Liquid Chromatography-Tandem Mass Spectrometry (LC-MS).

We replicated the Y-maze experiments and conducted an Object Recognition Test (ORT) in a new group of mice treated with either AAV-hSLC6A8 or PBS at P40 and P100. These tests were conducted at the CNR site (Baroncelli lab, Pisa). This study involved four groups of male mice: Slc6a8-KO (n=15) and WT (n=17) receiving i.c.v. injections of AAV-hSLC6A8, and Slc6a8-KO (n=18) and WT (n=16) receiving PBS. At P100, a subset of animals (n=4 per group) underwent post-mortem creatine (Cr) quantifications, which at the CNR site were carried out using Gas Chromatography-Mass Spectrometry (GC-MS). GC-MS Cr measurements were also performed at P20 and P40 in separate groups of Slc6a8-KO (n=4) and WT (n=4) animals treated with AAV-hSLC6A8, along with corresponding control groups of Slc6a8-KO (n=4) and WT (n=4) animals.

Finally, an additional group of animals composed of Slc6a8-KO (n=4) and WT (n=4) mice treated with AAV-hSLC6A8, and Slc6a8-KO (n=2), and WT (n=2) mice receiving PBS as control, were sacrificed at P40 for analysis of transgene biodistribution and cellular targeting via immunohistochemistry.

### AAV injection in newborn mice

Intracerebroventricular injection of AAV was performed as previously described^18^. Briefly, at P1 pups were placed on ice and then moved onto a cooled stereotaxic frame. Coordinates were adjusted on the base of lambda point of *repere* (X, Y, Z) = (1, ±0.3, - 2.0) mm. Pups received bilateral injections of 1ul of AAV vector (3x10^9^ vg/mouse). The AAV solution was infused over a 60s period and the pipette was removed after a delay of 30s to prevent backflow.

### Functional and anatomical magnetic resonance imaging (MRI)

Resting state fMRI (rsfMRI) data were acquired as described in^14,19–21^. Briefly, mice were anesthetized with isoflurane intubated and artificially ventilated. During resting-state scans, isoflurane was replaced with halothane (0.7%) to obtain light sedation. To assess potential differences in the sensitivity of Slc6a8-KO animals to the anesthetic used, we measured the Minimal Alveolar Concentration (MAC), in an independent group of adult mice, n=10 WT, n=5 Slc6a8-KO, as previously described^22,23^. We used a 7T MRI scanner (Bruker), Bruker Paravision software version 6, a 72mm birdcage transmit coil and a 4 channel solenoid coil for signal reception^22,24,25^. For each session, in vivo structural images were acquired with a fast spin-echo sequence. BOLD rsfMRI time series were acquired using an echo planar imaging (EPI) for 1920 volumes (total duration 32 minutes).

### Analysis of fMRI timeseries

rsfMRI BOLD time series were preprocessed as previously described^14,22,24,26^. Briefly, BOLD timeseries were despiked, motion corrected and spatially registered to a common reference brain template. Mean ventricular signal and motion traces of head realignment parameters were regressed out from time series. Finally, we applied a spatial smoothing and a band-pass filter to a frequency window of 0.01-0.1 Hz.

Network based statistics (NBS)^14,27^ was carried out by extracting fMRI signal in 85 anatomically parcellated regions (Table S2). The list of employed regions can be found in^28^. We next computed an unpaired two-tailed Student’s t test for each element of the corresponding correlation matrix separately (t>2.7-3.5). FWER correction at the network level was performed using 5000 random permutations (p<0.05). The chord plots show the 85 parcels clustered in 9 anatomical meta regions (Table S2). rsfMRI connectivity was also probed using seed-based analyses^24,29^. A seed region was selected to cover the areas of interest, based on NBS results (Supplementary Fig. 1). Voxel wise intergroup differences in seed-based mapping were assessed using a 2 tailed Student’s t test |t>2, p<0.05) and family wise error (FWE) cluster corrected using a cluster threshold of p=0.050.

### Brain volume analysis

T2-weighted anatomical scans were manually skull-stripped using ITKSNAP. We used the nipype python library^31^ wrapper for ants^32^ registration to compute symmetric diffeomorphic transform^33^ from dsurqe^34^ reference template on individual images. The obtained transform was applied on cortex, hippocampus, striatum, thalamus, using anatomical coordinates derived from the Paxinos/dsurqe ontology^35^.

### Immunohistochemistry

Conventional (single label) immunohistochemistry assays were carried out as described in supplementary methods. For double labeling experiments, brain sections were co-incubated overnight with primary antibodies against the HA tag, and targeting specific cell-type markers, including neuronal marker MAP2, astroglial marker GFAP, microglial marker Iba-1, oligodendroglial marker BCAS1. Antigen-antibody interactions were visualized using Alexa Fluor-conjugated secondary antibodies (1:400, Invitrogen).

### Quantification of HA-Tag Biodistribution and analysis of co-localization

The right hemisphere of each section was imaged using a Zeiss laser-scanning Apotome microscope. Maximum intensity projections (MIPs) were generated from the four consecutive optical sections displaying the highest mean pixel intensity. A threshold was established based on negative control HA-tag staining from PBS-injected animals. Subsequently, the area fraction of HA-positive pixels in the whole hemisphere was quantified.

For more targeted analysis, regions of interest (ROIs) were drawn around specific areas and the area fraction of HA-positive pixels within these ROIs was measured.

### Behavioral tests

To investigate spontaneous self-grooming, mice were individually placed in an open field arena for 20 minutes. Sessions were recorded and mice automatically tracked using the ANY-maze software. After a 10-min habituation period, the cumulative time spent by mice grooming themselves was scored for 10min as an index of stereotypic behavior^7,36^. Spatial working memory was tested using the Y-maze, as previously described^7,15^.

We also assessed declarative memory using the object recognition test (ORT). The total movement of the animal (Total exploration time) and the total distance covered (Distance moved) were automatically computed. A discrimination index was computed as previously reported^15^.

### Creatine measurements

#### Liquid Chromatography-Tandem Mass Spectrometry (LC-MS, performed at IIT)

Brain samples were homogenized in PBS-Protease inhibitor (100:1) on ice. An aliquot of each brain homogenate was extracted with cold CH_3_CN containing creatine- (methyl-*d3*) as internal standard and centrifuged at 21.100 x g for 20 min at 4°C. A calibration curve was prepared in PBS containing 20% CH_3_CN. The calibrators were extracted as the brain homogenates. The supernatants of the extracted brain homogenates and calibrators were further diluted 100-fold with 2mM NH_4_OAc in H2O (pH8) and analyzed by LC-MS/MS on a Waters ACQUITY UPLC-MS/MS system. Electrospray ionization was applied in positive mode. Compound-dependent parameters as Multiple Reaction Monitoring (MRM) transitions and collision energy were developed for the parent compound and the internal standard. The analyses were run on an ACE Excel 2 C18 (150x2.1mmID) with an ACE Excel UHPLC Pre-column Filter at 40°C, using 2mM NH4OAc in H_2_O (A) and CH3CN (B) as mobile phase at 0.2mL/min. All samples were quantified by MRM peak area response factor in order to determine the levels of Cr in the brain samples. Data were expressed as ng Cr per mg brain (ng/mg brain) and normalized by WT Creatine content (%WT).

#### Gas Chromatography-Mass Spectrometry (GC-MS, performed at CNR)

HEK293T pellets and brain samples were homogenized in 0.7mL of ice-cold PBS buffer using an ultrasonic disruptor. Samples were centrifuged at 600×g for 10 min at 4°C. A 50µl aliquot of the surnatant was taken for protein content assessment, while the remaining volume was used for Cr analysis, as previously described^7,37^. Data were expressed as nmol Cr per mg protein (nmol/mg pr) and normalized relative to the creatine content measured in wild-type (WT) samples (% WT).

### Partial Least Square Correlation

Partial-least square analyses were designed to maximize the covariance between imaging readouts (seed-based connectivity maps of the anterior cingulate), behavioral scores (i.e. self-grooming and spontaneous alternations) and brain Cr levels via the identification of latent components (LC) representing the optimal weighted linear combination of the original variables^38^. The null hypothesis that the observed brain-behavior relationship could be due to chance was verified via permutation testing (1,000 iterations) of behavioral and biochemical data matrix. Permutations were performed within groups as per previous guidelines^39^. The stability of the contribution of each brain and behavioral element was assessed via bootstrapping, where bootstrap resampling was performed within each experimental group to avoid patterns being driven by group differences^40^.

## Data availability

All the imaging data will be made publicly available to researchers upon acceptance for publication of our study.

## Results

### Creatine deficiency disrupts fMRI connectivity in cortical and subcortical brain regions

To probe whether CTD affects interareal functional connectivity, we carried out resting-state fMRI (rsfMRI) measurements in Slc6a8-KO mice and control littermates (WT) at two different stages of disease progression. Specifically, we longitudinally mapped fMRI connectivity in the same mice at postnatal day 40 (P40) and in adulthood (P140), corresponding to early and late pathological stage, respectively. Animals were scanned using an established sedation protocol that has been shown to preserve network organization in rodents, making it comparable to that mapped in awake conditions^41,42^. Moreover, the protocol has proven effective in identifying genotype-dependent effect in multiple genetic models of developmental disorders^43^, revealing highly congruent functional connectivity changes in corresponding clinical populations^21,22,44^. To obtain a spatially-unbiased mapping of genotype-specific differences in fMRI connectivity we used Network Based Statistics (NBS) using a whole-brain network parcellation^14,27^. This analysis revealed the presence of diffuse hypoconnectivity in multiple brain regions of Slc6a8-KO mice both at P40 and P140 (Fig. 1). The spatial extent and magnitude of the observed changes, however, were largely different across the probed ages. In juvenile mice, fMRI hypoconnectivity was distributed across multiple cortical and subcortical regions, with a prominent involvement of somatosensory and motor cortices as well as hippocampal and thalamic areas (Fig. 1a,b). In contrast, fMRI hypoconnectivity in Slc6a8-KO mice at P140 was weaker and more focal, encompassing the motor cortex and subcortical regions such as the striatum, the hypothalamus and some thalamic areas (Fig. 1c,d). To further investigate the circuit-level substrates differentially affected by CTD, we next used a seed-based analysis to probe fMRI connectivity in some of the brain regions exhibiting large effect-size in NBS (Fig. 1). This investigation revealed robustly reduced inter- hemispheric connectivity in somatomotor areas of Slc6a8-KO mice at P40 (Fig. 2a-d), and to a lower extent also at P140 (Fig. 2e,h; Supplementary Fig. 1). At P40, Slc6a8-KO mice also showed reduced thalamo-hippocampal and frontocortical connectivity, but no alterations in striatal regions (Supplementary Fig. 2). In contrast, the striatum represented one of the primary hubs of fMRI hypoconnectivity in Slc6a8-KO mice at P140, with evidence of decreased coupling of this region with thalamic nuclei (Fig. 2f,h). Adult Slc6a8-KO mice also exhibited foci of reduced fMRI connectivity in thalamo-hippocampal regions (Supplementary Fig. 2). Importantly, these connectivity differences are unlikely to reflect a different sensitivity of Slc6a8-KO animals to the employed sedative regime, because Minimal Alveolar Concentration (MAC) testing in a separate group of animals revealed highly comparable sensitivity to halothane across the two genotypes (WT, MAC 1.70 ± 0.13; Slc6a8-KO, MAC 1.68 ± 0.08; student t test p = 0.87). Leveraging the longitudinal design of our investigation, we also examined the temporal evolution of connectivity in somato-motor and striatal circuits across the two timepoints tested (Supplementary Fig. 3). A pairwise quantification of connectivity in these networks did not reveal any difference in temporal trajectories across groups (Supplementary Fig. 3). Altogether, these findings reveal widespread disruption of brain connectivity in Slc6a8-KO mice, with robust alterations of somato-motor connectivity in juvenile mice evolving into more focal cortical and subcortical hypoconnectivity in adulthood.

**Figure 1.**
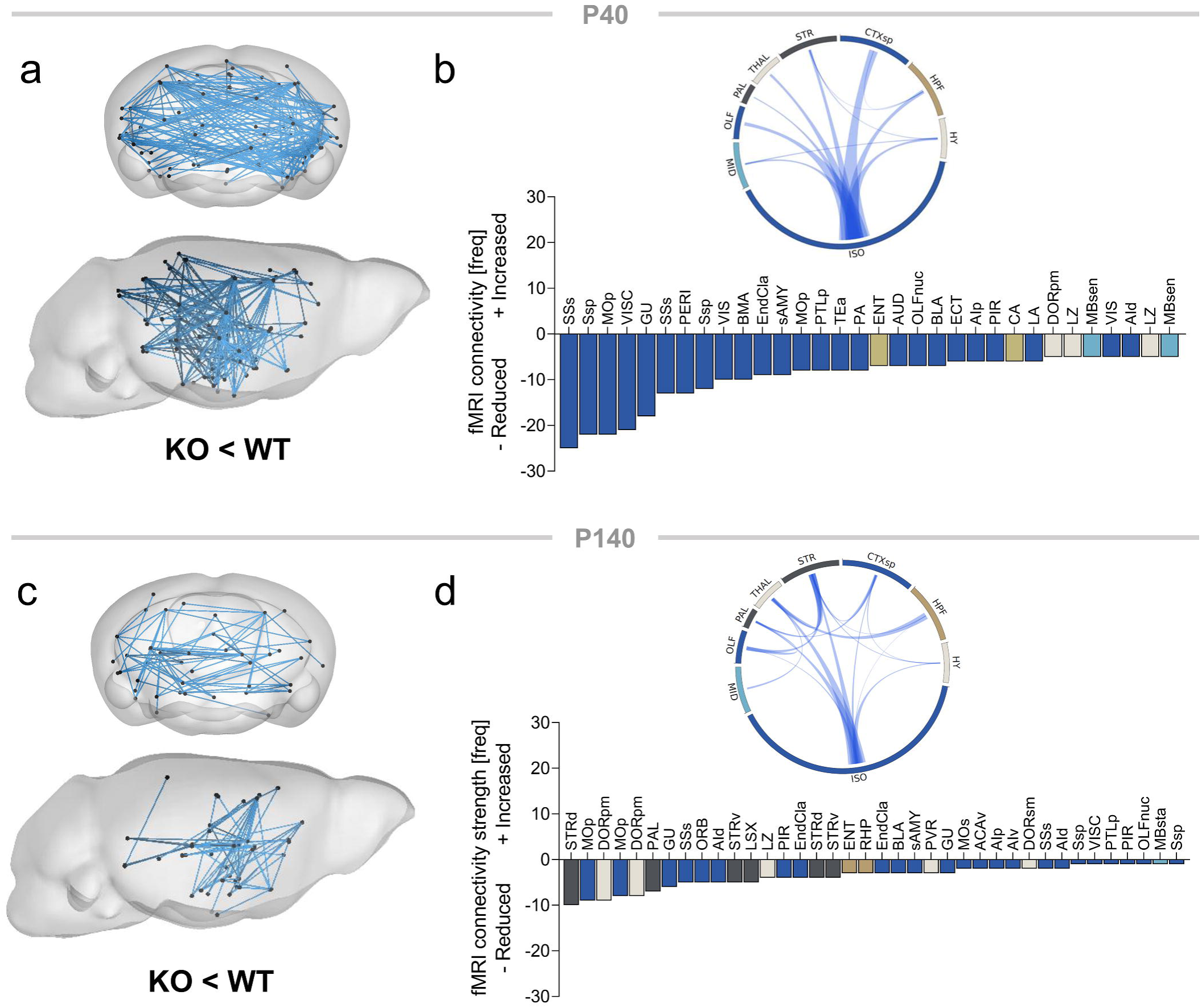
Reduced long-range fMRI connectivity in Slc6a8-KO mice. fMRI connectivity alterations in Slc6a8-KO mice as assessed with NBS at P40 (**a, b**) and P140 (**c, d**; t> |2.7| for both comparisons). Histograms represent the number of links exhibiting reduced connectivity in Slc6a8-KO mice at P40 and P140, respectively. Circular plots in (**b**) and (**d**) illustrate corresponding patterns of altered interareal connectivity. In these plots, individual brain areas have been grouped into 9 anatomical meta regions. Thickness of links in circular plots is proportional to the relative number of affected links. The complete list of brain areas (and abbreviations) used for NBS is reported in Table S2.

**Figure 2.**
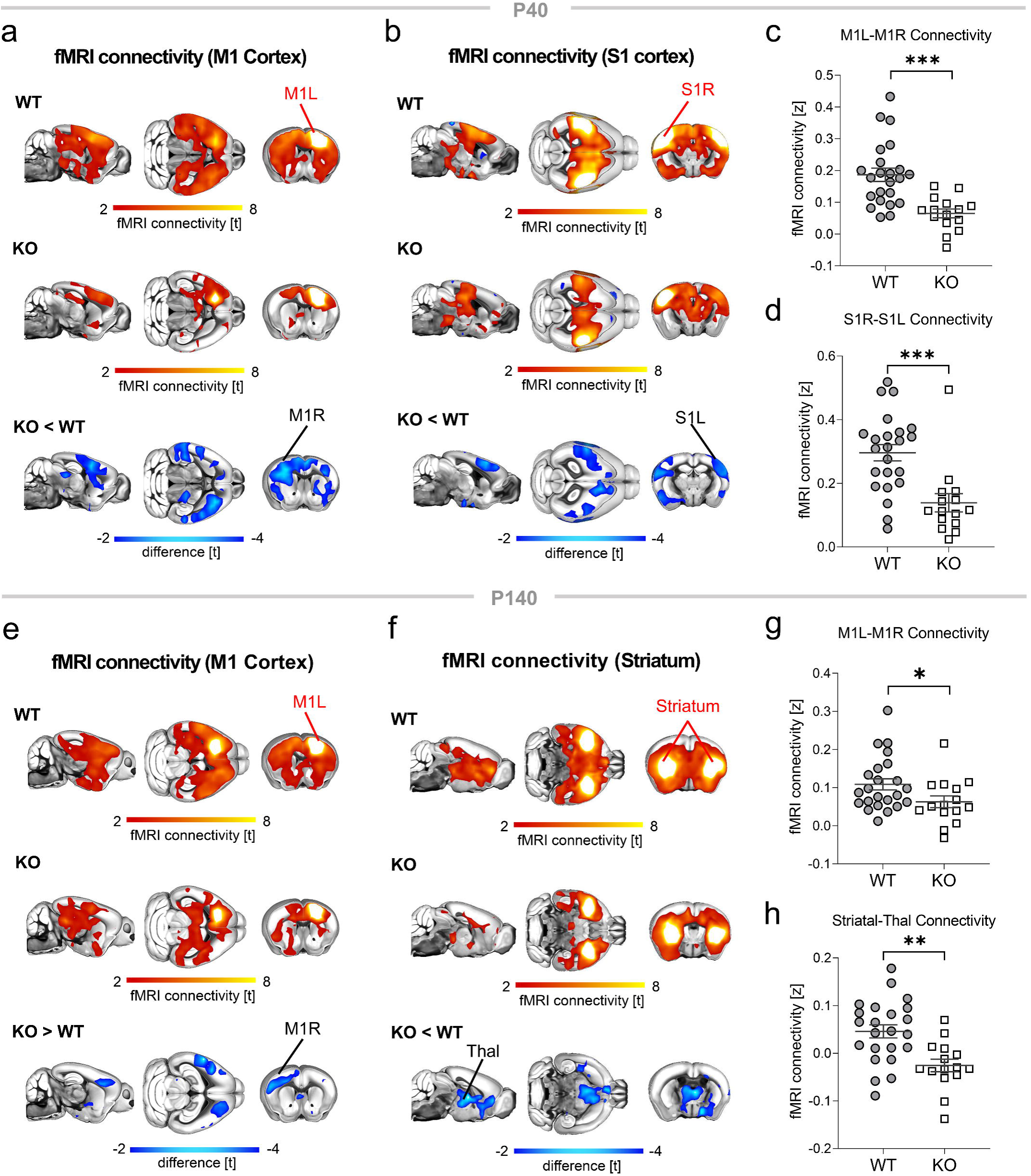
Functional networks exhibiting altered fMRI connectivity in Slc6a8-KO mice. (**a-d**) Seed-correlation mapping highlighted disrupted somato-motor connectivity in Slc6a8-KO mice at P40. (**e-h**) fMRI connectivity in motor regions was disrupted also at P140. At this age we also found a prominent reduction of fMRI connectivity between striatal regions and the thalamus. Red/yellow indicates areas in the brain maps exhibiting significant (t > 2.1) fMRI connectivity with seed regions (indicated with red lettering). Blue indicates between-group connectivity differences (t-test, t > 2.1). Corresponding quantifications of connectivity changes in the two groups are reported in panels c,d for P40 (S1 t = 4.02, p = 0.0003; M1 t = 4.3, p = 0.0001) and g,h for P140 (M1, t = 2.06, p = 0.046; Striatal-Thal, t = 3.55, p =0.001). L, left; R, right; M1, motor cortex; S1, somatosensory cortex; Thal, thalamus. All statistics are FWE cluster-corrected. FWE, family-wise error. *p < 0.05, **p < 0.01, ***p < 0.001. Error bars indicate SEM and dots represent individual values.

### AAV-mediated expression of human SLC6A8 prevents fMRI hypoconnectivity in juvenile Slc6a8-KO mice

To probe the efficacy of genetic expression of exogenous CRT as putative therapy for CTD, we conducted intracerebroventricular (i.c.v.) injections of a novel AAV9-vector encoding a functional copy of the non-codon optimized human *SLC6A8* gene under the control of JeT promoter (AAV9/JeT-hSLC6A8, herein referred to as AAV-hSLC6A8). The AAV-hSLC6A8 vector was designed to leverage the wide biodistribution of AAV following i.c.v. perinatal injection^45^ and to allow broad and ubiquitous transgene expression in the brain via the use of JeT promoter^46^. Our experimental goal was to induce early hSLC6A8 expression to minimize or prevent the emergence of pathological CTD-related phenotypes. Before proceeding with *in vivo* efficacy testing, we conducted a series of control studies to probe the function, transduction efficiency, and tropism of the employed vector.

We first verified that the HA-tagged version of CRT we employed would not affect its function. To this end, we compared Cr uptake in 293T cells transfected with either the original plasmid encoding the HA-tagged transporter, or a newly developed plasmid encoding the native (untagged) transporter. Notably, we observed highly comparable Cr uptake between the two conditions (204.00 ± 13.32 and 208.00 ± 16.64 nmol/mg protein for the tagged and untagged version, respectively, one-way ANOVA, post hoc Tukey’s test, p = 0.97), suggesting that the incorporation of an HA-tag in the hSLC6A8 gene does not impair CRT function.

We next mapped *in vivo* transduction patterns of exogenous CRT using an anti-HA-tag antibody (Fig. 3a, Supplementary Fig. 4a). We found that transgenic CRT was expressed in a punctate pattern along cell membranes as expected. The transduction pattern of exogenous CRT exhibited widespread extension in both WT and Slc6a8-KO mice (Fig. 3a, Supplementary Fig. 4a). Expression was particularly prominent in the deeper layers of the cortex and extended to subcortical regions, including the hippocampus and the striatum. Quantitative analyses indicated an overall transduction efficiency of 7.7% across experimental groups (7.6 ± 3.7% and 7.8 ± 1.3% in Slc6a8-KO and WT animals, respectively; Fig. 3b, Supplementary Fig. 4b). Transduction efficiency was not affected by genotype (two-way ANOVA, genotype effect, p= 0.37) but exhibited regional variability, with lower efficiency in deep parenchymal regions (e.g. striatum, two-way ANOVA, region effect, p= 0.01). To better define the cellular targets of hSLC6A8 gene transfer and determine whether transduction differentially affects neurons and glial cells, we next performed double-labeling of HA-tag with cell-type-specific markers, including MAP2 for neurons, GFAP for astrocytes, IBA1 for microglia, and BCAS1 for oligodendrocytes. Our analysis demonstrated a strong tropism of the AAV-hSLC6A8 vector for neurons (Fig. 3c and Supplementary Fig. 4c), as well as oligodendrocytes and microglia, whereas no astrocytes were found to be double-labeled with the HA tag (Supplementary Fig. 5-7).

**Figure 3.**
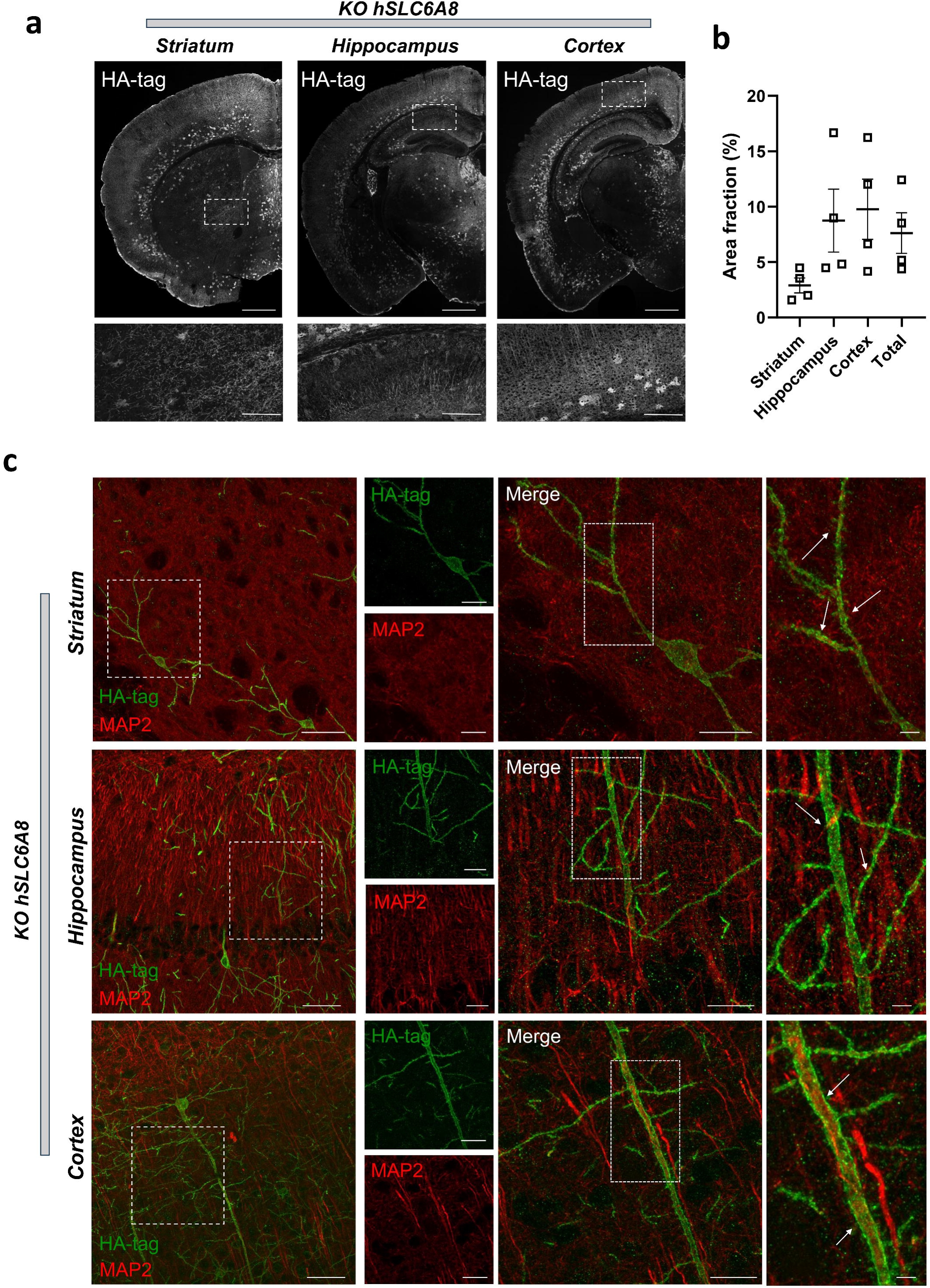
Exogenous CRT protein is widely distributed throughout the brain and colocalizes with neurons in different brain regions in Slc6a8-KO mice. (**a**) Representative coronal sections (20x magnification) stained to quantify HA-tagged distribution of transgenic CRT protein in the striatum, hippocampus and cerebral cortex of Slc6a8-KO mice treated with AAV-hSLC6A8. Scale bars are 1 mm top panels, 100 µm bottom panels. (**b**) Quantification of the area positive for HA-tag staining (%) in the three considered brain regions, and in the right brain hemisphere (total) of Slc6a8-KO mice. Error bars indicate SEM. (**c**) Representative images illustrating colocalization of HA-tag staining (green signal) along with the neuronal marker MAP2 (red signal) across different brain regions. Panels on the leftmost column were acquired at 20x magnification (scale bars are 50 µm). Additional panels were acquired at 63x magnification (scale bars are 20 µm, 50 µm and 5 µm from left to right, respectively).

To test whether perinatal AAV-mediated expression of hSLC6A8 represents a viable gene therapy for CTD, we carried out longitudinal fMRI and behavioral investigation. The AAV-hSLC6A8 vector was injected perinatally in newborn (P1) Slc6a8-KO mice and WT littermates (Fig. 4a). An analogous vector expressing GFP protein (AAV-GFP) was used as control condition in both genotypes. Notably, NBS analysis of fMRI measurements revealed that AAV-hSLC6A8 administration prevented the manifestation of fMRI hypoconnectivity in Slc6a8-KO mice at P40 (Fig. 4b,c). This effect entailed a marked increase of fMRI connectivity in virtually all the neocortical and subcortical regions that we found to be hypo-connected in Slc6a8-KO mice injected with AAV-GFP. Seed-based mapping corroborated this result, showing that gene therapy could effectively ameliorate somato-motor (Fig. 4d-g), and cingulate hypoconnectivity in Slc6a8-KO mice at P40 (Supplementary Fig. 8). In contrast, no appreciable connectivity normalization was observed in adulthood in the same mice, using NBS analysis (p > 0.9, t < 0.1, all links). Accordingly, seed-based mapping did not reveal any functionally relevant effect in treated Slc6a8-KO mice at P140 (Fig. 4h-k; Supplementary Fig. 8). Taken together, these results show that neonatal expression of hSLC6A8 robustly improved juvenile fMRI hypoconnectivity in Slc6a8-KO mice, but not the corresponding alterations in fMRI connectivity observed at a later pathological stage (i.e., P140).

**Figure 4.**
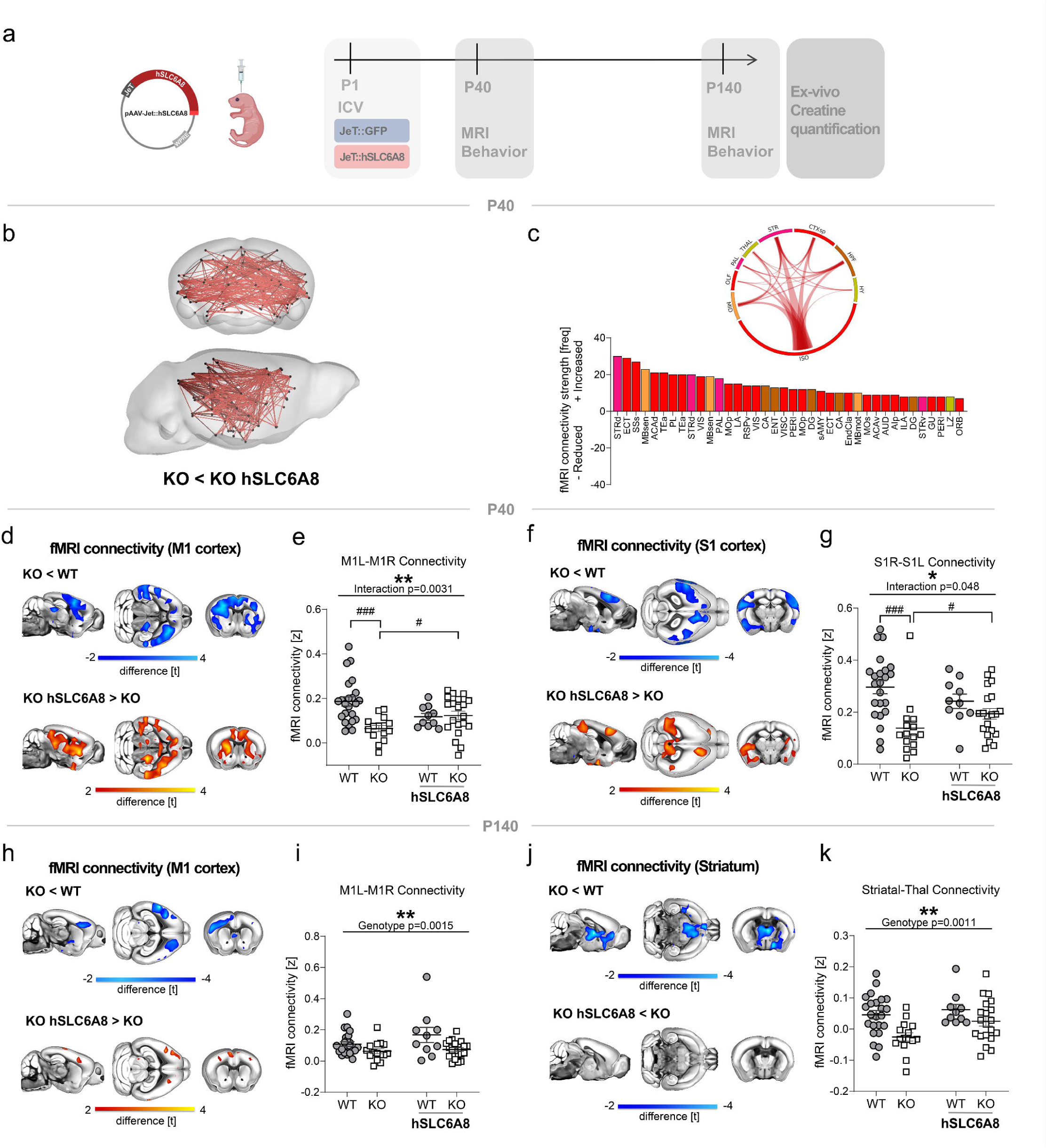
Perinatal AAV-hSLC6A8 injection prevents onset of fMRI hypoconnectivity in juvenile Slc6a8-KO mice. (**a**) Experimental design. (**b**) NBS highlighted increased fMRI connectivity at P40 in AAV-treated Slc6a8-KO mice compared to controls (red, t > |3|). (**c**) Connectivity links exhibiting a significant amelioration of connectivity in AAV-treated Slc6a8-KO mice and regional quantification in circular plots. (**d-g**) Seed-correlation mapping revealed increased interhemispheric connectivity in the somato-motor cortex of AAV-treated juvenile Slc6a8-KO mice. Red/yellow coloring in bottom panels denotes the amelioration of fMRI connectivity produced by AAV-hSLC6A8 treatment (t-test, t > 2). Plots report quantification of connectivity strength between seed and regions exhibiting amelioration of connectivity (**e**, M1: two-way ANOVA, genotype x treatment interaction F = 9.43, p = 0.0031; t-test, # p < 0.05; ### p < 0.001; **g**, S1: two-way ANOVA, interaction F = 4.06, p = 0.048; Mann Whitney test # p < 0.05, t-test, ### p < 0.001). The same analysis at P140 revealed no effect of gene therapy in either motor cortex (**i**; two-way ANOVA, genotype factor F = 11.05, p = 0.0015, interaction F =1.30, p = 0.26) or the striatum-thalamus network (**k**; two-way ANOVA, genotype factor F = 11.77, p = 0.0011, interaction F = 1.16, p = 0.28). L, left; R, right; S1, somatosensory cortex; M1, motor cortex; Thal, thalamus. All statistics are FWE cluster-corrected. FWE, family-wise error. For two-way ANOVA, * p< 0.05, **p< 0.01, ***p < 0.001. Error bars indicate SEM and dots represent individual values. The complete list of brain areas (and abbreviations) used for NBS is reported in Table S2.

We finally tested whether our gene therapy would affect brain anatomical alterations in Slc6a8-KO mice. To this aim, we analyzed longitudinal anatomical T2- weighted images acquired from the same mice used for fMRI at P40 and P140. Whole-brain volumetric analysis revealed a small but highly significant reduction in brain volume in Slc6a8-KO mice at both ages (P40: Δ = -4.9%; P140: Δ = -5.9%; Supplementary Fig. 9). Normalized regional volume quantifications showed no significant differences in all of the examined regions (thalamus, cortex, striatum, hippocampus) at P40 and P140. Importantly, AAV-mediated hSLC6A8 expression did not affect these volumetric reductions at either age (Supplementary Fig. 9). These findings reveal a generalized reduction in brain volume in Slc6a8-KO animals, which remained unaffected by gene therapy.

### AAV-mediated expression of human SLC6A8 prevents onset of autistic-like behavior in SLC6A8-KO mice

We previously reported that Slc6a8-KO mice exhibit a number of traits and phenotypes of translational relevance for CTD, including reduced body weight, cognitive impairment and autistic-like behavior^7,12^. To assess whether perinatal gene therapy could prevent the onset of CTD-relevant pathological manifestations, we probed working memory (using the Y maze task) and stereotyped movements (via self-grooming scoring), and recorded body weight at P40 and P140 in mice injected with either AAV-hSLC6A8 or AAV-GFP (Fig. 5).

**Figure 5.**
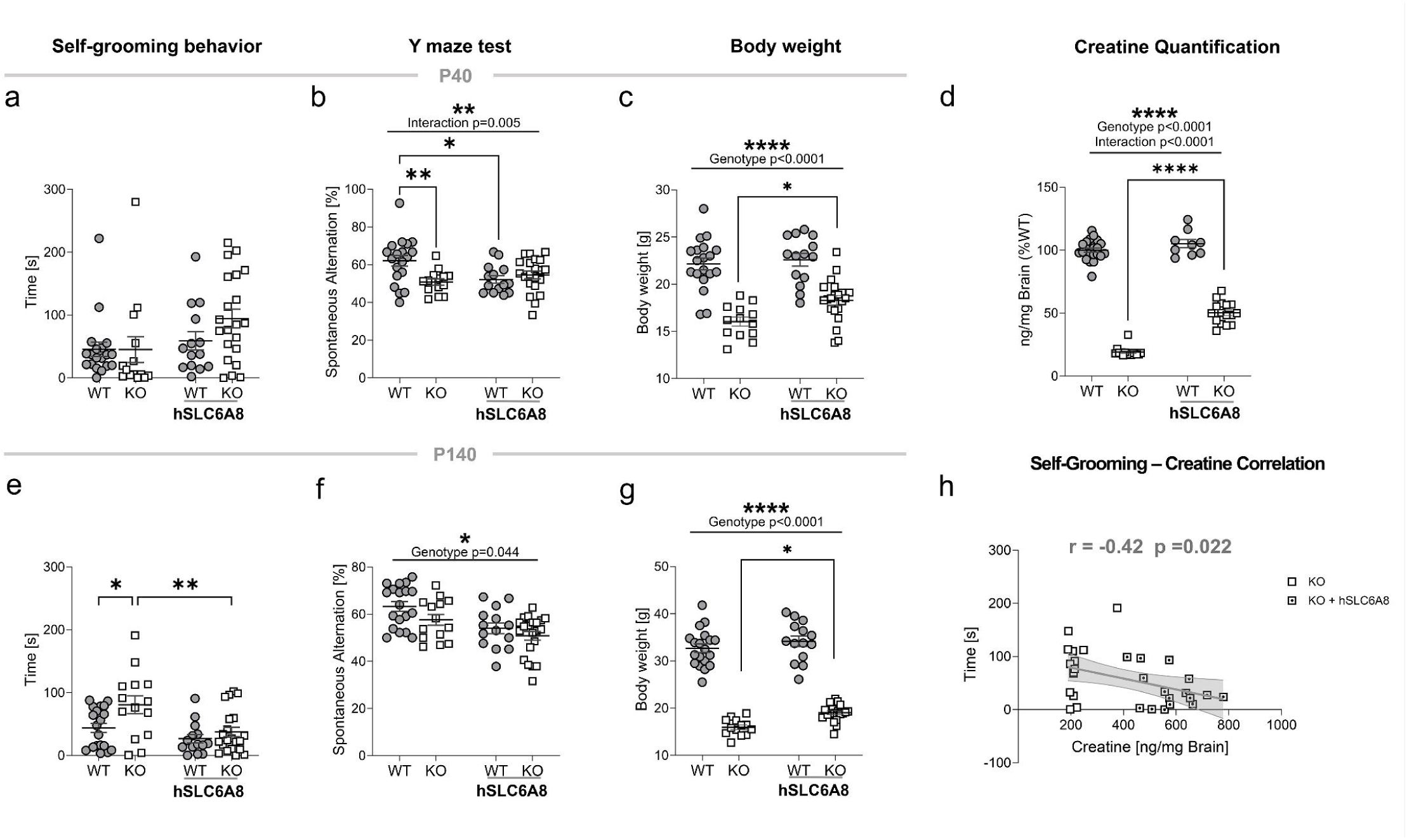
Perinatal AAV-hSLC6A8 injection increases body weight and prevents the onset of autistic-like stereotypies in adult Slc6a8-KO mice. (**a-c**) Time spent self-grooming (**a**, two-way ANOVA, genotype x treatment interaction F = 1.33, p = 0.25), spontaneous alternation in the Y maze task (**b**, two-way ANOVA, interaction F = 8.55, p = 0.005) and body weight (**c**, two-way ANOVA, genotype factor F = 77.81, p < 0.0001, interaction F = 2.32, p = 0.13) in the four experimental groups at P40. (**d**) Quantification of brain creatine levels at P140 (two-way ANOVA, genotype factor F = 1068, p < 0.0001, interaction F = 39.63, p < 0.0001). (**e**) AAV-hSLC6A8 injection prevented onset of aberrant grooming behavior in adult (P140) Slc6a8-KO mice (two-way ANOVA, interaction F = 2.09, p = 0.15), while no effect was found in the Y maze (**f**, two-way ANOVA, genotype factor F = 4.23, p = 0.044, interaction F = 0.34, p = 0.56). (**g**) Body weight of treated Slc6a8-KO mice was significantly increased compared to controls (two-way ANOVA, genotype factor F = 420, p < 0.0001, interaction F = 1.08, p = 0.30). (**h**) Brain creatine levels negatively correlate with self-grooming scoring in Slc6a8-KO mice (Pearson r = -0.42, R squared = 0.17, p = 0.022). This correlation was carried out using the same data in panel e. Note that the treated KO group includes fewer animals because some mice were randomly selected for immunohistochemical assessments to verify viral expression. Error bars indicate SEM and each dot represents a mouse. Tukey’s multiple comparison test, *p < 0.05, **p < 0.01, ***p < 0.001, ****p < 0.0001.

Slc6a8-KO mice at P40 did not show aberrant grooming behavior (Fig. 5a), but exhibited significantly decreased spontaneous alternation in the Y maze (Fig. 5b), and reduced body weight compared to WT mice (Fig. 5c). The same mice showed significantly increased grooming (Fig. 5e), decreased spontaneous alternations (Fig. 5f) and robustly reduced body weight at P140 (Fig. 5g), thus recapitulating our previous findings^7^.

Perinatal gene therapy produced a significant amelioration of some of these pathological traits. Specifically, at P40 we found a moderate (≈15%) increase of body weight of Slc6a8-KO mice, but no effect on behavioral performance in the spontaneous alternation test. Remarkably, at P140 AAV-hSLC6A8 treatment completely prevented the appearance of increased grooming in Slc6a8-KO mice. This effect was accompanied by a partial (≈20%) gain of body weight. However, as seen in juvenile mice, gene therapy failed to improve behavioral performance in the spontaneous alternation task at this disease stage. The inclusion of a group of WT mice receiving an AAV-hSLC6A8 injection allowed us also to investigate the behavioral effect of CRT overexpression. We noted a slight deterioration of behavioral performance in the Y maze task in WT mice receiving the AAV-hSLC6A8 vector (Fig. 5b,f), as well as increased baseline exploratory activity (i.e., number of arm entries) in both Slc6a8-KO and WT mice injected with AAV-hSLC6A8 (Supplementary Fig. 10).

To further strengthen our behavioral investigations, we replicated the Y-maze experiments, and conducted the Object Recognition Test (ORT) in a new group of mice treated with either AAV-hSLC6A8 or PBS at two distinct developmental time points (P40 and P100, Fig. 6). These tests were conducted at a separate site (CNR, Pisa) using an independent mouse colony, allowing us to assess the generalizability of our primary behavioral findings. Consistent with our previous results (obtained at IIT, Rovereto), perinatal AAV-hSLC6A8 treatment did not improve Y-maze performance in Slc6a8-KO mice at either time point (Fig. 6a,d). Additionally, we observed significantly reduced Y-maze performance in AAV-hSLC6A8-treated WT mice at P40, with a similar, though non-significant, trend towards poorer performance at P100 (Fig. 6a,d).

**Figure 6.**
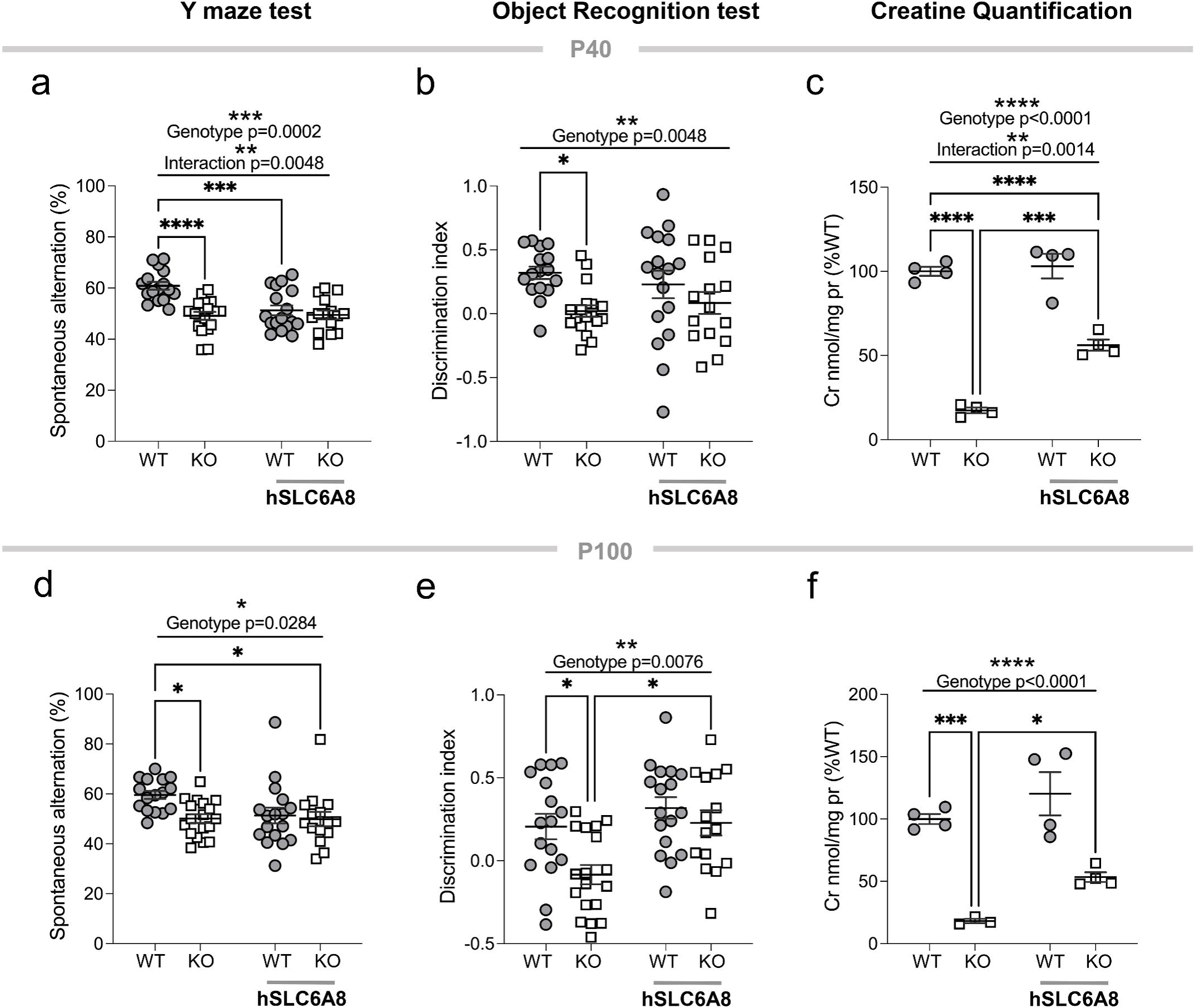
Perinatal AAV-hSLC6A8 injection partially restores Cr levels and cognitive function in adult Slc6a8-KO mice. (**a**) At P40, AAV-hSLC6A8 treatment impaired Y-maze performance in WT mice, while no changes were detected in Slc6a8- KO mice (two-way ANOVA, genotype factor F = 15.83, p = 0.0002, treatment F = 7.37, p = 0.0086, interaction F = 8.55, p = 0.0048). At the same age, no effect of gene therapy was observed in either WT or hSLC6A8-KO mice in the ORT (**b**, two-way ANOVA, genotype factor F = 8.58, p = 0.0048). (**c**) Quantification of brain creatine levels at P40 (two-way ANOVA, genotype factor F = 225, p < 0.0001, treatment F = 23.5, p = 0.0004, interaction F=17.0.9, p = 0.0014). (**d**) At P100, AAV-hSLC6A8 injection did not ameliorate behavioral performance in Slc6a8-KO mice in the Y-maze (two-way ANOVA, genotype factor F = 5.04, p = 0.0284), but significantly improved cognitive performance in the ORT (**e)**, two-way ANOVA, genotype factor F = 7.62, p = 0.0076, treatment F = 9.54, p = 0.003). (**f**) Quantification of brain creatine levels at P100 (genotype factor F = 56.03, p < 0.0001, treatment F = 7.81, p = 0.0174). Error bars indicate SEM and each dot represents a mouse. Tukey’s multiple comparison test, *p<0.05, **p<0.01, ***p<0.001, ****p<0.0001.

Importantly, AAV-hSLC6A8 administration significantly mitigated cognitive deficits of Slc6a8-KO mice in the ORT at P100 (discrimination index, Fig. 6e). No significant therapeutic effect was instead observed at P40 (Fig. 6b). Total exploratory behavior was comparably increased in both WT and Slc6a8-KO mice treated with AAV- hSLC6A8 at P100, thus ruling out genotype-specific confounding effects of general locomotor activity in the ORT (Supplementary Fig. 11). Taken together, these behavioral results demonstrate that perinatal gene therapy in Slc6a8-KO mice can alleviate specific pathological traits of high translational relevance for CTD, including cognitive dysfunction and autism-like motor stereotypies.

### Gene therapy with AAV-SLC6A8 increases brain creatine levels

Brain Cr depletion underlies the pathological cascade leading to CTD^2,3^. Consequently, the therapeutic effects of AAV-hSLC6A8 treatment we observed on fMRI connectivity, motor stereotypies, declarative memory and body weight should reflect higher brain Cr levels. To verify this, we measured post-mortem Cr levels in distinct groups of animals at P20 (Supplementary Fig. 12), P40 (Fig. 6c), and in a randomly selected subgroup of mice at the end of behavioral investigations at P100 and P140 (Fig. 5d and Fig. 6f). Our findings revealed a significant increase in brain Cr levels in AAV-hSLC6A8-treated KO mice. At the earliest time point (P20), Cr levels in Slc6a8-KO mice reached approximately 70% of WT values (Supplementary Fig. 12). This increase stabilized at around 50–55% of WT levels by P40, and remained stable throughout the study (Fig. 5d, Fig. 6c,f). These results indicate that AAV-mediated expression hSLC6A8 encodes significant amounts of functionally-active CRT protein. It should be noted that the employed gene therapy was not able to fully reinstate physiological levels of Cr in Slc6a8-KO mice, as the average Cr content measured in mutant mice receiving AAV-hSLC6A8 was approximately half the amount measured in WT mice. In spite of this, we found an inverse relationship between Cr brain levels and self-grooming scoring in Slc6a8-KO mice treated with AAV-GFP or AAV-hSLC6A8 (Fig. 5h; Pearson r = -0.42; p = 0.022). A similar inverse relationship and slope was also noticeable in Slc6a8-KO mice receiving AAV-hSLC6A8 injections, although the effect in this group did not reach statistical significance (Fig. 5h; Pearson r = -0.35; p = 0.18). These results suggest that perinatal AAV-hSLC6A8 administration can effectively increase central levels of Cr, resulting in domain-specific amelioration of CTD pathology.

### Connectivity of a mesolimbic-prefrontal circuit predicts behavioral performance, and is modulated by creatine levels

To obtain an unbiased mapping of the putative circuits underlying behavioral disruption in our CTD mouse model, we used a multivariate model to associate cross-subject variance in fMRI connectivity with corresponding behavioral profiles. This was done using a partial least square correlation (PLS) analysis^38–40,47^ on fMRI connectivity data extracted from the cingulate cortex. The use of multivariate approaches like PLS reduces the bias related to the use of univariate brain/behavior correlations in relatively small samples, an approach that has been recently shown to be highly prone to false positives^48^. Another advantage of PLS is the possibility to model, within the same framework, Cr levels as a continuous variable. Thus, using PLS we sought to identify areas whose connectivity covaries with (and thus putatively explains) the behavioral profile across multiple tests.

For each animal we modeled behavioral scores out of spontaneous alternation and grooming tests, and Cr levels measured at P140 (Fig 7). By including Cr levels as a continuous variable, we were able to assess whether the identified circuits are sensitive to Cr modulation. We probed connectivity of the cingulate cortex because this region projects to multiple brain areas that are relevant to the tests examined, including ventral hippocampus for spontaneous alternation, and striatal and mesolimbic areas for self-grooming^22,49^.

**Figure 7.**
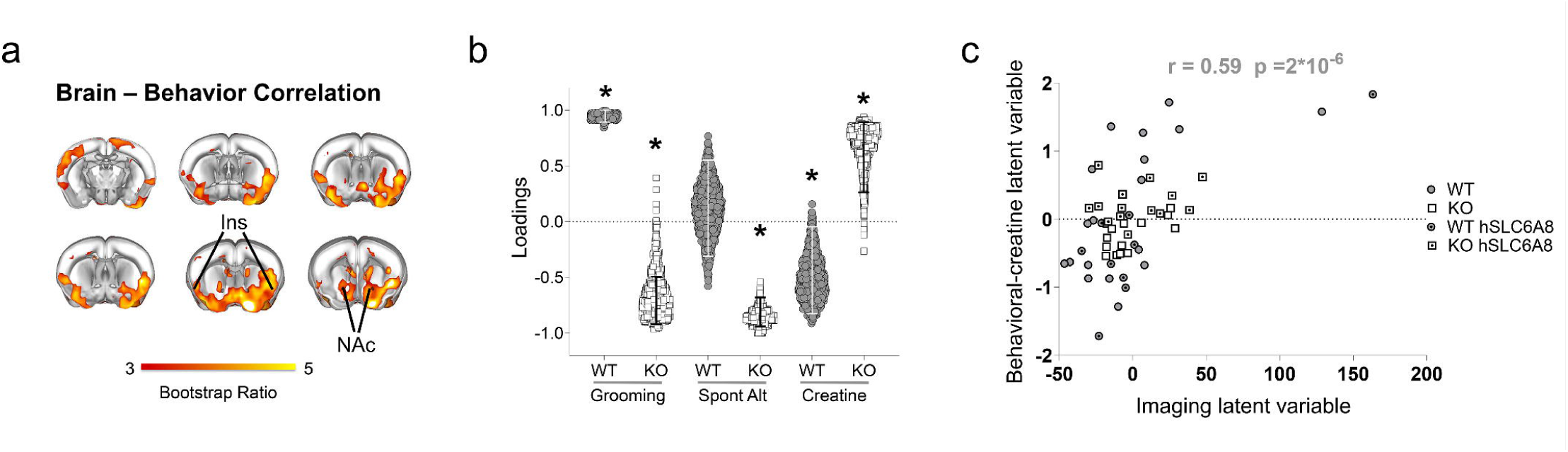
Multivariate modeling reveals disrupted brain-behavior relationship in Slc6a8-KO mice. PLS analysis revealed a significant latent component (LC1) that explained 40% of covariance between self-grooming, cingulate-striatal-insular network and Cr levels across mice. **a**) Red/yellow coloring represents bootstrap-ratio (BSR) z-scores obtained from 1000 bootstrapping iterations. The highlighted basal-forebrain and mesolimbic regions are those whose connectivity with the anterior cingulate exhibit robust covariance with behavioral performance and Cr levels as per the employed PLS model. **b**) Creatine and behavior loadings for LC1. Columns indicate the contribution of each observed variable to the LC and the directionality of covariance. Errors bars indicate 5^th^ to 95^th^ bootstrapping percentiles. Asterisk indicates a significant contribution to the overall PLS relationship for LC1 (relationship is non-zero, i.e., 95% CIs do not encompass 0 loading). **c**) PLS correlation between individual brain and behavioral- creatine scores for LC1, where different groups are represented by dot shape and color. The Behavioral-Creatine PLS variable indicates a latent dimension that captures shared variance between self-grooming, spontaneous alternation measures and Cr levels. Each dot indicates an individual mouse. Ins, Insular Cortex; NAc, Nucleus Accumbens.

PLS analysis revealed one highly significant latent component (LC1; p = 0.002) accounting for 40% of the covariance between brain connectivity, behavioral performance and Cr levels. Regional mapping of the identified component revealed a prefrontal-basal-forebrain (i.e., insula, nucleus accumbens) circuit whose connectivity positively covaried with stereotyped behavior, but not spontaneous alternation, in WT mice (Fig 7a,b). This result is consistent with the known organization of the distributed network of regions that control stereotyped activity in rodents^49^ and suggest that connectivity measurements we employed are behaviorally relevant. Interestingly, this brain-behavior relationship was disrupted in Slc6a8-KO mice, where we found reversed covariance between behavioral scores and connectivity in the same network.

Importantly, Cr levels were also found to covary (and thus potentially modulate) the activity of this circuit, albeit with different directional effects in WT and Slc6a8-KO mice (Fig 5b,c). In control mice, Cr levels were inversely related to the activity in these regions, suggesting that non-homeostatic levels of Cr can negatively affect this circuit to impact behavior. In contrast, Cr levels were found to be positively related to behavioral performance in Slc6a8-KO mice, a finding consistent with the ameliorating effect of AAV-mediated hSLC6A8 expression.

These results uncover a behaviorally-relevant prefrontal-basal-forebrain network whose activity is differentially modulated by Cr levels. This brain-behavior relationship was disrupted in Slc6a8-KO mice, hence implicating aberrant prefrontal-mesolimbic connectivity in the pathology of CTD.

## Discussion

Recent advancements in understanding the clinical manifestations and etiopathological course of CTD^2,3,12^ have not been paralleled by a deeper comprehension of the systems-level dysfunction and mechanisms underlying this genetic syndrome. To fill this knowledge gap, we have employed longitudinal functional imaging, behavioral phenotyping and genetic therapy in an established murine model of CTD to investigate how Cr depletion affects brain connectivity and leads to the circuit dysfunctions that characterize CTD. By using longitudinal mapping, we were able to track the longitudinal evolution of fMRI dysconnectivity in the same animal group, with minimal invasiveness, and high precision. Our fMRI studies revealed robust functional hypoconnectivity both in juvenile and adult Slc6a8-KO mice, suggesting that Cr depletion leads to impaired large-scale interareal communication. We found fMRI hypoconnectivity to mostly affect cortical regions during early pathological stages, subsequently evolving into more prominent subcortical hypoconnectivity (especially in striatal, mesolimbic and thalamic areas) in adulthood. Although fMRI hypoconnectivity may reflect multifactorial mechanisms^50^, our results are consistent with prior evidence of disrupted function and maintenance of synaptic circuits in Slc6a8-KO mice^8^. Synaptic coupling is a key determinant of long-range synchronization underlying fMRI connectivity, and recent investigations have revealed fMRI hyperconnectivity in autism-relevant mouse models exhibiting aberrant synaptic signaling^22^, as well as fMRI hypoconnectivity in mouse models showing reduced synaptic density^13,14^. In this respect, our results corroborate the notion that brain connectopathy is a hallmark endophenotype of multiple neurodevelopmental disorders associated with synaptic pathology^22,43^.

The observation of fMRI hypoconnectivity in Slc6a8-KO mice is also in agreement with our previous study showing that these mice exhibit a severe epileptic phenotype and significant changes in neural oscillations, with lower power of theta/alpha EEG frequencies and increased power of beta/gamma bands^9^. Chemogenetic and pharmacological manipulations have indeed shown that fMRI hypoconnectivity can reflect reduced low frequency EEG power and concomitant broad- band increased in higher-frequency activity as a result of increased excitability and asynchronous firing^24,29^. Thus, the effect of Cr deficiency on EEG power aligns with the observed reduction in fMRI connectivity (i.e., a measure of infraslow synchronization), potentially indicating a shift from slow, synchronized neural coherency to high frequency (yet asynchronous) activity. Similar spectral changes are present in the EEG of CTD children with respect to age-matched controls^9^, underscoring the translational value of the present study.

From a cellular standpoint, these alterations may be linked to prior studies pointing at a significant heterogeneity of *Slc6a8* expression across cell populations^5,51,52^, and the observation that synaptic alterations in CTD mostly affect GABAergic interneurons^7,8^. These alterations may in turn lead to a generalized reduction of cortical inhibitory tone and a disruption of neural circuitry efficiency, two key pathological determinants associated with multiple neurodevelopmental disorders^53^. Given the crucial role of parvalbuminergic (PV) fast-spiking interneurons in the regulation of long-range functional synchronization^54^, the presence of morpho-physiological dysfunction of PV synapses in Slc6a8-KO mice^8^ might represent a plausible cellular correlate of the observed fMRI desynchronization. Accordingly, previous studies have shown that alterations of the excitatory/inhibitory ratio may perturb brain oscillatory activity and fMRI connectivity with potential specific contributions of different cell populations^14,29^.

The putative translational relevance of these imaging findings is further corroborated by the observation of decreased inter-hemispheric connectivity in Slc6a8- KO mice at both P40 and P140. Given the robust anatomical foundations of functional connectivity^28^ and the causal role of the corpus callosum in driving the synchronization of the two brain hemispheres^55–57^, these findings may be functionally linked to previous anatomical MRI studies revealing corpus callosum thinning in CTD patients^12^.

Our study also serves as a preclinical investigation of the therapeutic potential of early gene therapy in preventing the pathological and behavioral manifestations of CTD. The chronic progressive morbidity of CTD represents a significant unmet clinical need for which gene therapy could potentially offer a lifelong treatment option. In this respect, CTD is an ideal target for gene additive therapy for three reasons: i) it is a monogenic condition; ii) the replenishment of brain Cr is effective in ameliorating the clinical manifestations of two other disorders caused by alterations in Cr metabolism^58,59^; iii) the expression of functional Slc6a8 and the severity of CTD phenotypes exhibit a dose-dependent relationship, with heterozygous females for *Slc6a8* mutations showing moderate Cr reduction and partial cognitive deficits^60,61^. Interestingly, intracisternal delivery of an AAV2/9 vector carrying the rat Slc6a8 gene under a CMV promoter (targeting the cerebellum, medulla oblongata and spinal cord) prevented myocytic and locomotor impairments in a rat model of CTD but failed to improve other behavioral domains (e.g. cognition, or stereotyped movements)^62^. This result adds to the present findings to suggest that key etiopathological phenotypes can be partly rescued by viral-mediated gene therapy in CTD.

In the present study, a single intraventricular infusion of AAV-hSLC6A8 in newborn Slc6a8-KO mice was sufficient to achieve widespread expression of the transgene across multiple brain regions. The observed transduction is broadly consistent with previously reported efficiency of AAV9-mediated vectors following intracerebroventricular administration^63,64^, resulting in a sustained and stable increase in cerebral Cr levels. Importantly, although Cr concentration in Slc6a8-KO brains remained approximately 50% lower than physiological levels measured in WT mice, postnatal reinstatement of CRT function led to partial of full rescue of key CTD-relevant phenotypes. These included functional brain hypoconnectivity in juvenile mice, reduced body weight, impaired declarative memory and the manifestation of autistic-like stereotyped behaviors in adulthood. The independent assessment of cognitive functions in two separate laboratories strengthens the robustness and reproducibility of these behavioral findings.

Interestingly, our PLS analysis linked these endophenotypes to the activity of a fronto-mesolimbic network whose function was disrupted in Slc6a8-KO mice, thus revealing a neural circuitry potentially central to CTD pathology, and whose function was modulated by Cr levels in adulthood. These findings are consistent with previous studies showing that reinstatement of cerebral Cr metabolism via oral Cr supplementation can mitigate clinical symptoms in other neurodevelopmental syndromes associated with mutations of Cr synthetic enzymes^65–67^. However, a limitation of our PLS approach is that the identified latent variable represents a statistical combination of distinct behavioural measures rather than a single coherent behavioural domain. Future studies incorporating additional behavioral assessments are required to further refine the brain-behavior associations reported here.

Rather unexpectedly, AAV-mediated delivery of hSLC6A8 did not fully restore cognitive performance in Slc6a8-KO mice, with significant improvements only observed in the ORT at P100. Several factors may account for this finding. Since optimal AAV expression requires about two weeks, our perinatal injection strategy might have failed to provide physiologically-relevant Cr levels during an early time-window of critical importance for the maturation of higher order cognitive functions^68^. Additionally, the viral titer employed in our study might not have reached the transduction threshold sufficient to fully prevent the onset of CTD pathological endophenotypes. Indeed, creatine levels in AAV-hSLC6A8-treated Slc6a8-KO mice remained well below the physiological concentrations observed in WT controls, suggesting that complete restoration of physiological creatine concentrations may be necessary to achieve normal cognitive functioning in CTD. In keeping with this, we recently reported a strictly dose-dependent therapeutic effect of cyclocreatine (a lipophilic creatine analogue capable of substituting its metabolic function) on cognitive defects and stereotyped behavior in Slc6a8-KO mice, possibly reflecting distinct metabolic constraints of the different underlying brain processes^65^. Moreover, we cannot exclude the possibility that different cellular populations may require distinct physiological Cr levels for optimal function (e.g. PV interneurons^8^). If this is the case, ubiquitous transgene expression driven by JeT promoter might result in non-optimal (supraphysiological) expression of CRT in some cell populations, and insufficient expression in others. This might also explain the partial impairment of mnemonic performance observed in AAV-hSLC6A8-treated WT mice at P40, an effect consistent with a potential detrimental effect of non-homeostatic CRT expression levels.

Finally, the discrepancy between the lack of efficacy in the Y-maze and the partial improvement observed in the ORT may reflect differing metabolic Cr demands across brain regions. Given that declarative memory primarily relies on prefrontal and perirhinal cortical circuits^69^, whereas the Y-maze probes hippocampal-dependent spatial working memory^70^, hippocampal circuits might require complete full Cr replenishment to achieve full functionality. In contrast, prefrontal and cortical circuits may retain partial efficacy at lower Cr concentrations. These requirements could interact with, or be entirely driven by, differences in the maturation timeline of these circuits^71,72^. Further preclinical studies are warranted to optimize CRT expression such to achieve physiological replenishment of Cr levels in the central nervous system of Slc6a8-KO mice.

We finally note that a limitation of this study is the lack of electrophysiological analyses of the epileptic phenotype. Due to the fragility of KO animals and the relative tardive onset of the epileptic phenotype, in our study we prioritized earlier timepoints and longitudinal behavioral studies with the aim to enable direct brain-behavior correlations.

In conclusion, our results document translationally relevant, systems-level disruption of brain activity in a murine model of CTD and provide key proof-of-concept evidence that early gene therapy holds potential as a disease modifying strategy or CTD. We expect these findings to help the development of experimental therapies for this severe genetic disorder.

## Supplementary material

Supplementary material is available.

