## Supplementary Information for "Connectomic and behavioral alterations in creatine transporter deficiency are partially normalized by gene therapy"

**Author affiliations:**

[§] Contributed equally

[+] Contributed equally

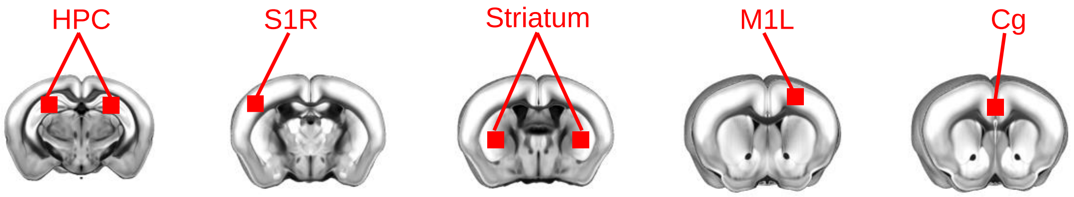


***Supplementary Figure 1. Anatomical location of seed regions.*** *HPC, hippocampus; S1R, right somatosensory cortex 1; M1L, left motor cortex; Cg, cingulate cortex.*

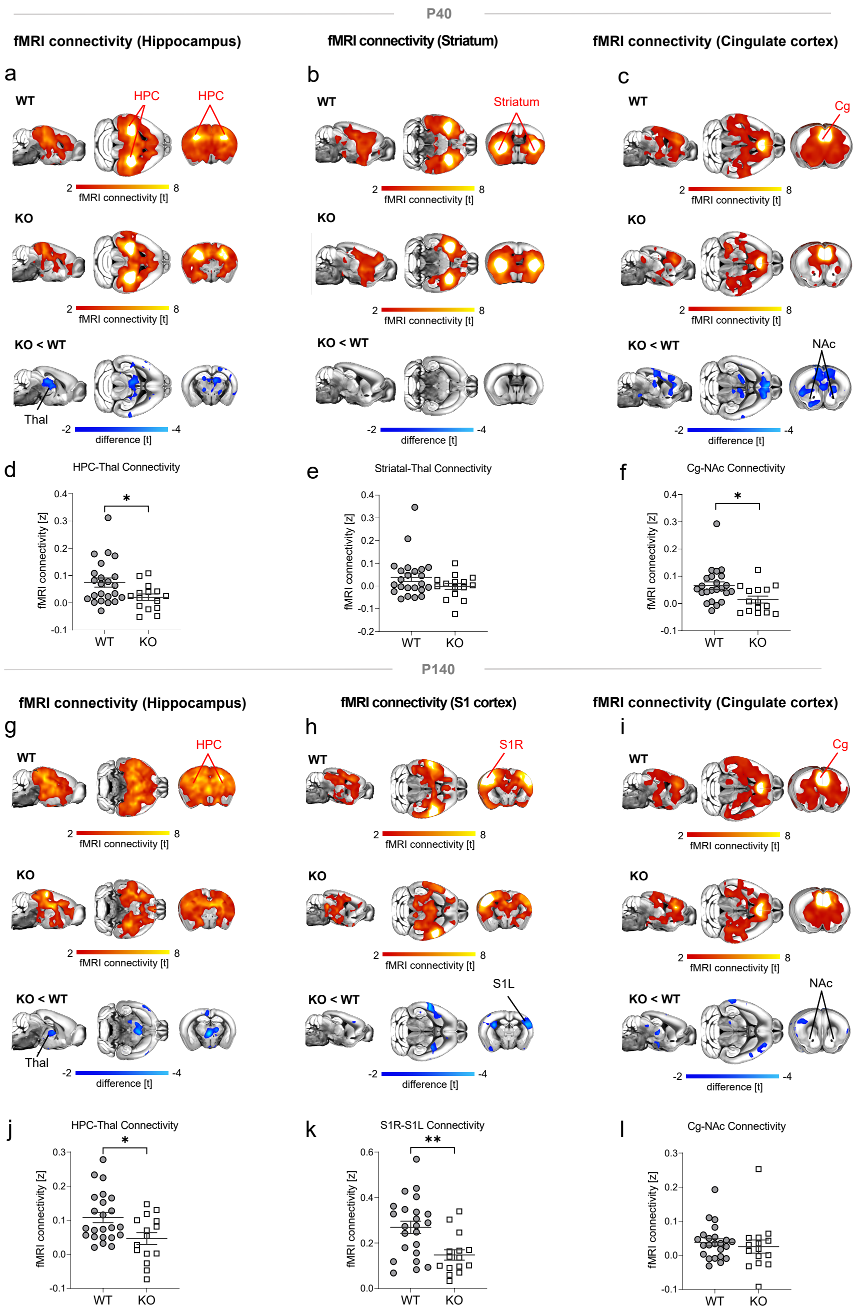


***Supplementary Figure 2. Seed-based analysis of fMRI connectivity.*** *(a-f) Seed-correlation mapping highlighted disrupted hippocampal-thalamus and cingulate cortex connectivity in Slc6a8-KO mice at P40. (g-l) fMRI connectivity in the hippocampal circuit was disrupted also at P140. At this age we also found a prominent reduction of inter-hemispheric somatosensory connectivity. Red/ yellow indicates areas in the brain maps exhibiting significant (t> 2.1) fMRI connectivity with seed regions (indicated with red lettering). Blue indicates between-group connectivity differences (t-test, t> 2.1). Corresponding quantifications of connectivity changes in the two groups are reported in panels d-f for P40 (HPC-Thal, t-test, t= 2.35, p= 0.025; Cg-NAc, t-test; t= 2.6, p= 0.014) and j-l for P140 (HPC-Thal, t-test; t= 2.65, p= 0.012; S1R-S1L, t-test; t = 3.17, p = 0.003). HPC, hippocampus; Thal, thalamus; Cg, cingulate cortex; NAc, Nucleus Accumbens; S1R, right somatosensory cortex; S1L, left somatosensory cortex. All statistics are FWE cluster-corrected. FWE, family-wise error. *p < 0.05, **p < 0.01. Error bars indicate SEM and dots represent individual values.*

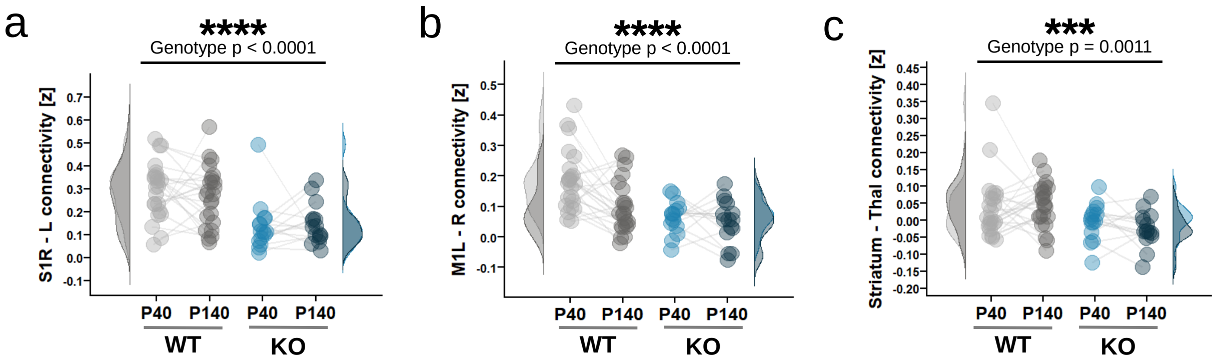


***Supplementary Figure 3. Temporal evolution of fMRI connectivity in WT and Slc6a8-KO mice at P40 and P140.*** *Quantification and distribution of fMRI connectivity in somatosensory (a, two-way RM ANOVA, genotype factor F = 26, p < 0.0001, genotype x age interaction F = 0.43, p = 0.512) and motor cortices (b, genotype factor F = 14.2, p = 0.0003, genotype x age interaction F = 2.8, p = 0.1) and between striatal and thalamic regions (c, genotype factor, F = 11.4, p = 0.0012, interaction F = 0.85, p = 0.36) in WT and Slc6a8-KO mice administered with AAV-GFP, at P40 and P140. ***p < 0.001, ****p<0.0001. L, left; R, right; M1, motor cortex; S1, somatosensory cortex; Thal, thalamus. Dots represent individual values.*

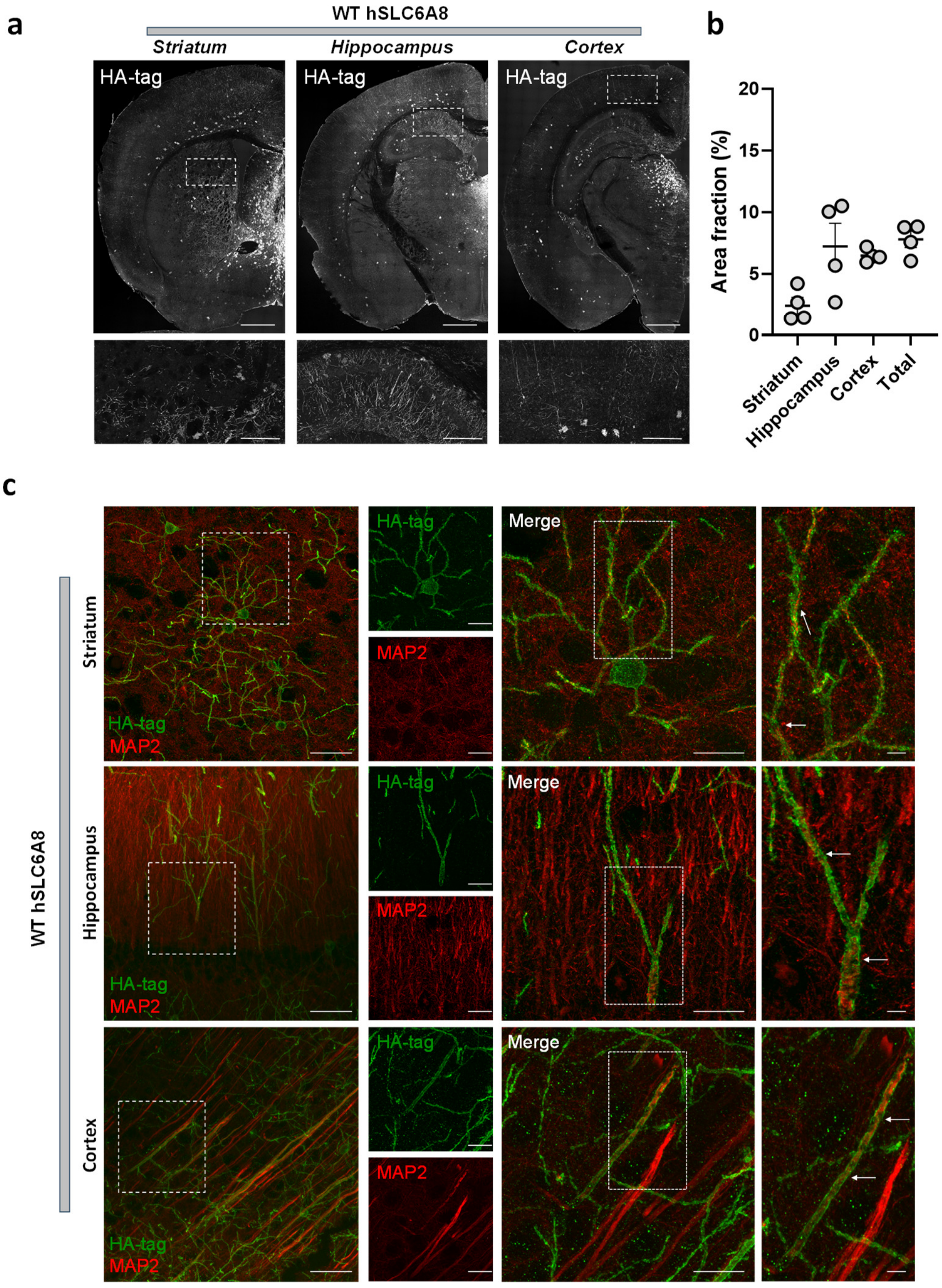


***Supplementary Figure 4. Exogenous CRT protein is widely distributed throughout the brain and colocalizes with neurons in different brain regions in WT mice.*** *(**a**) Representative coronal sections (20x magnification) stained to quantify HA-tagged distribution of transgenic CRT protein in the striatum, hippocampus and cerebral cortex of WT mice treated with AAV-hSLC6A8. Scale bar are 1 mm top panels, 100 µm bottom panels. (**b**) Quantification of the area positive for HA-tag staining (%) in the three brain regions considered and in the right brain hemisphere (total) of WT mice. Error bars indicate SEM. (**c**) Representative images illustrating colocalization of HA-tag staining (green signal) along with the neuronal marker MAP2 (red signal) across different brain regions. Panels on the leftmost column were acquired at 20x magnification (scale bars are 50 µm). Additional panels were acquired at 63x magnification (scale bars are 20 µm, 50 µm and 5 µm from left to right, respectively).*

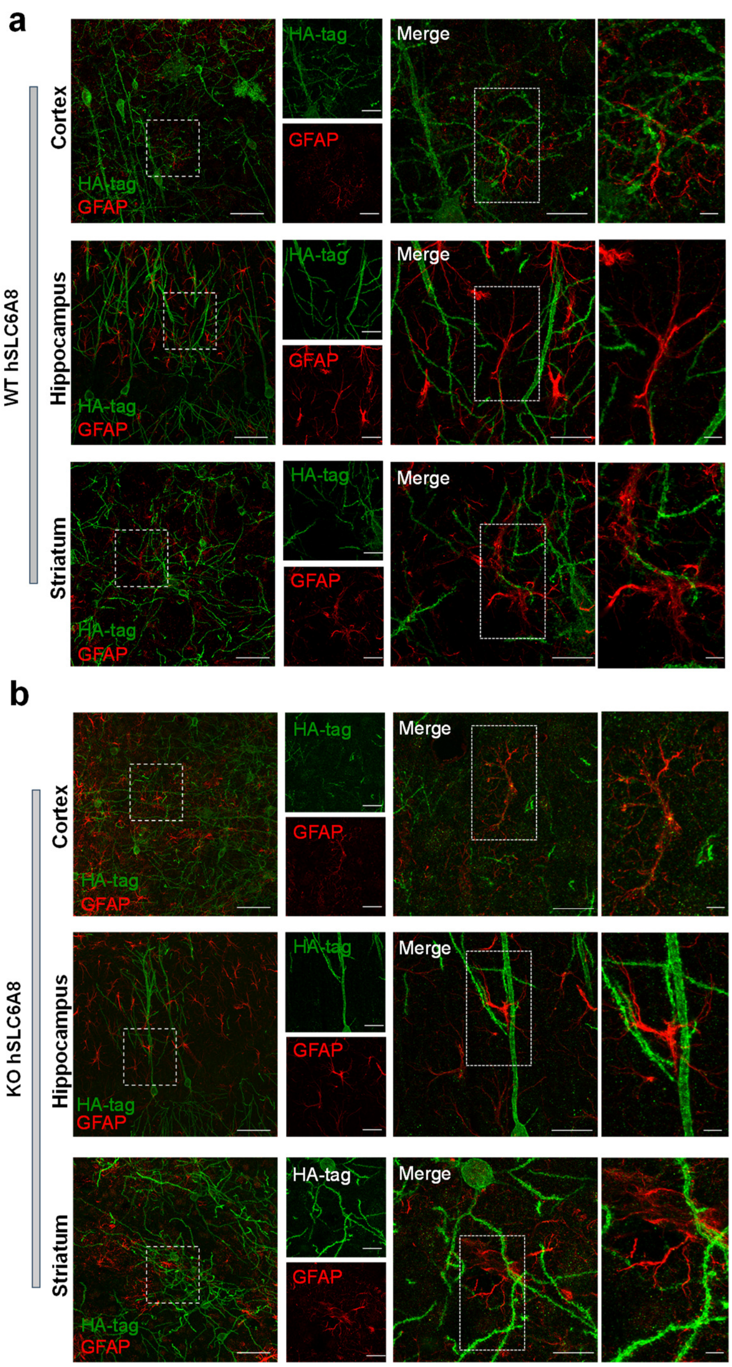


***Supplementary Figure 5. Exogenous CRT protein is not expressed in astrocytes.*** *Representative images illustrating colocalization of HA-tag staining (green signal) with the astrocytic marker GFAP (red signal) in the cortex, hippocampus and striatum of WT* ***(a)*** *and Slc6a8-KO* ***(b)*** *mice treated injected with AAV-hSLC6A8. In both (a) and (b), images in the leftmost columns were acquired at 20x magnification (scale bars are 50 µm). Additional panels were acquired at 63x magnification (scale bars are 20 µm, 50 µm and 5 µm from left to right, respectively).*

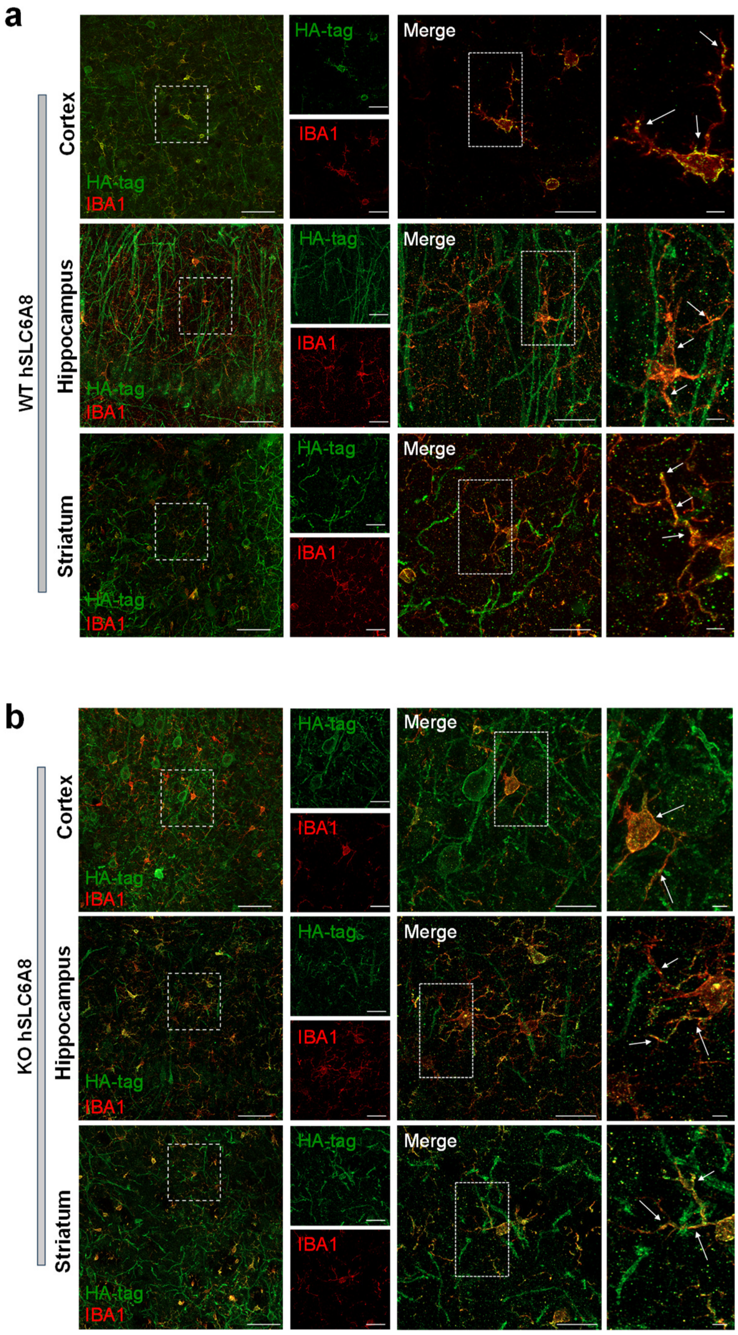


***Supplementary Figure 6. Colocalization of exogenous CRT protein with the microglial marker IBA1.*** *Representative images at 20x and 63x magnification, and relative insets illustrating the colocalization of HA-tag staining (green signal) with the microglial marker IBA1 (red signal) in the cortex, hippocampus and striatum of WT **(a)** and Slc6a8-KO **(b)** mice injected with AAV-hSLC6A8 at P1. In both (a) and (b), images in the leftmost columns were acquired at 20x magnification (scale bars are 50 µm). Additional panels were acquired at 63x magnification (scale bars are 20 µm, 50 µm and 5 µm from left to right, respectively).*

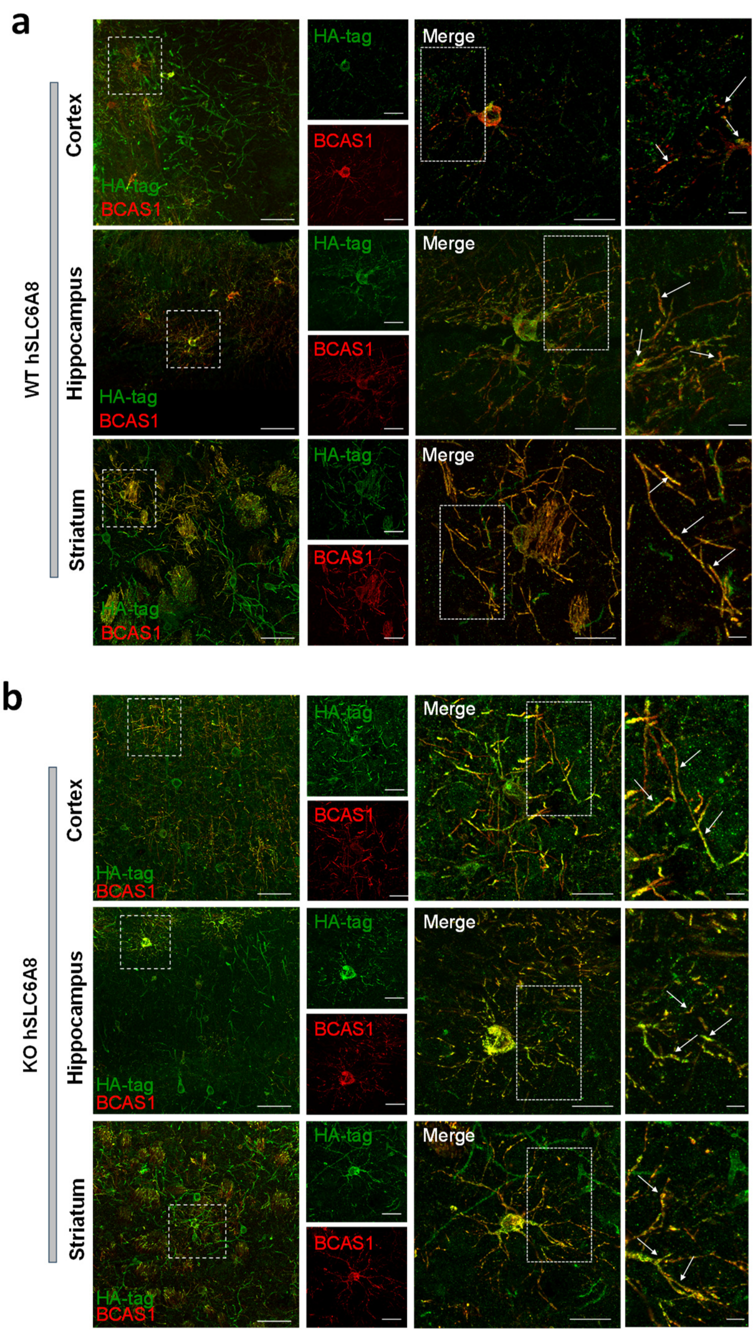


***Supplementary Figure** 7. **Exogenous CRT protein is expressed in oligodendrocytes.** Representative images at 20x and 63x magnification, and relative insets illustrating the colocalization of HA-tag staining (green signal) with oligodendrocytic marker BCAS1 (red signal) in the cortex, hippocampus and striatum of WT **(a)** and Slc6a8-KO **(b)** mice injected with AAV-hSLC6A8. In both (a) and (b), images in the leftmost columns were acquired at 20x magnification (scale bars are 50 µm). Additional panels were acquired at 63x magnification (scale bars are 20 µm, 50 µm and 5 µm from left to right, respectively).*

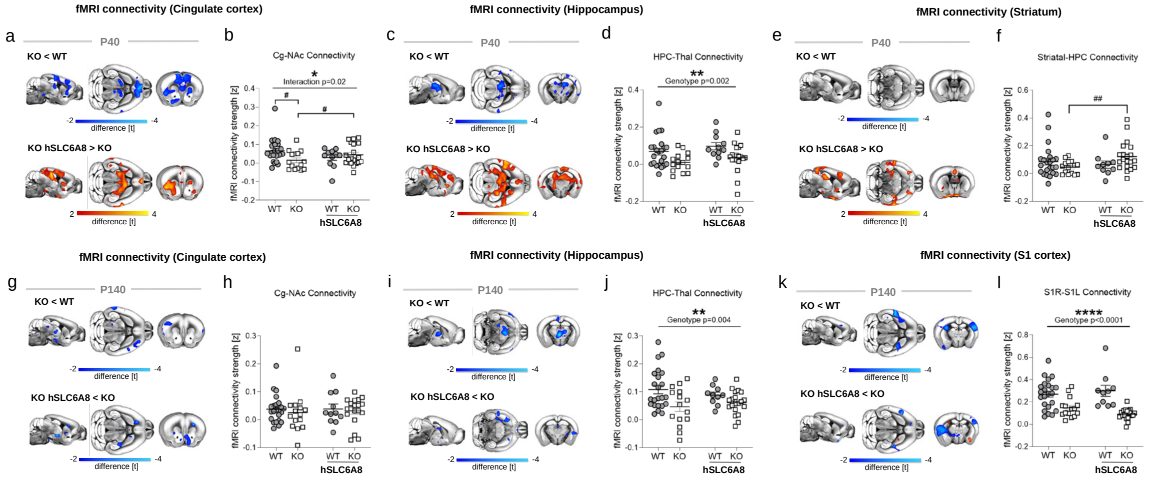


***Supplementary Figure 8. AAV-hSLC6A8 treatment prevents onset of fMRI hypoconnectivity in Slc6a8-KO mice.*** ***(a-f)*** *Seed-correlation mapping highlighted ameliorated fronto-limbic hypoconnectivity in treated Slc6a8-KO mice at P40. Red/yellow coloring in bottom panels denotes increased connectivity produced by AAV- hSLC6A8 treatment (t-test, $t > 2$). Histograms illustrate quantification of connectivity strength between the probed seed and regions of interest (b, Cg-NAc: two-way ANOVA, genotype x treatment interaction $F = 5.24$, $p = 0.025$; t-test, # $p < 0.05$; d, HPC-Thal: two-way ANOVA, genotype factor $F = 10.7$, $p = 0.0017$, interaction, $F = 0.01$, $p = 0.92$; f, striatal-HPC: two-way ANOVA, interaction $F = 3.94$, $p = 0.051$; t-test, ## $p < 0.01$).* ***(g-l)*** *The same analysis at P140 revealed no effect of gene therapy (h, Cg-NAc: two-way ANOVA, interaction $F = 0.03$, $p = 0.85$; j, HPC-Thal: two-way ANOVA, genotype factor $F = 8.77$, $p = 0.004$, interaction, $F = 1.33$, $p = 0.25$; l, S1R-S1L: two-way ANOVA, genotype factor $F = 36.5$, $p < 0.0001$, interaction $F = 2.60$, $p = 0.11$). HPC, hippocampus; Ins, insula; Thal, thalamus; Cg, cingulate cortex; NAc, Nucleus Accumbens; S1R, right somatosensory cortex 1; S1L, left somatosensory cortex 1. All statistics are FWE cluster-corrected. FWE, family-wise error. For two-way ANOVA, * $p < 0.05$, ** $p < 0.01$, **** $p < 0.0001$. Error bars indicate SEM and dots represent individual values.*

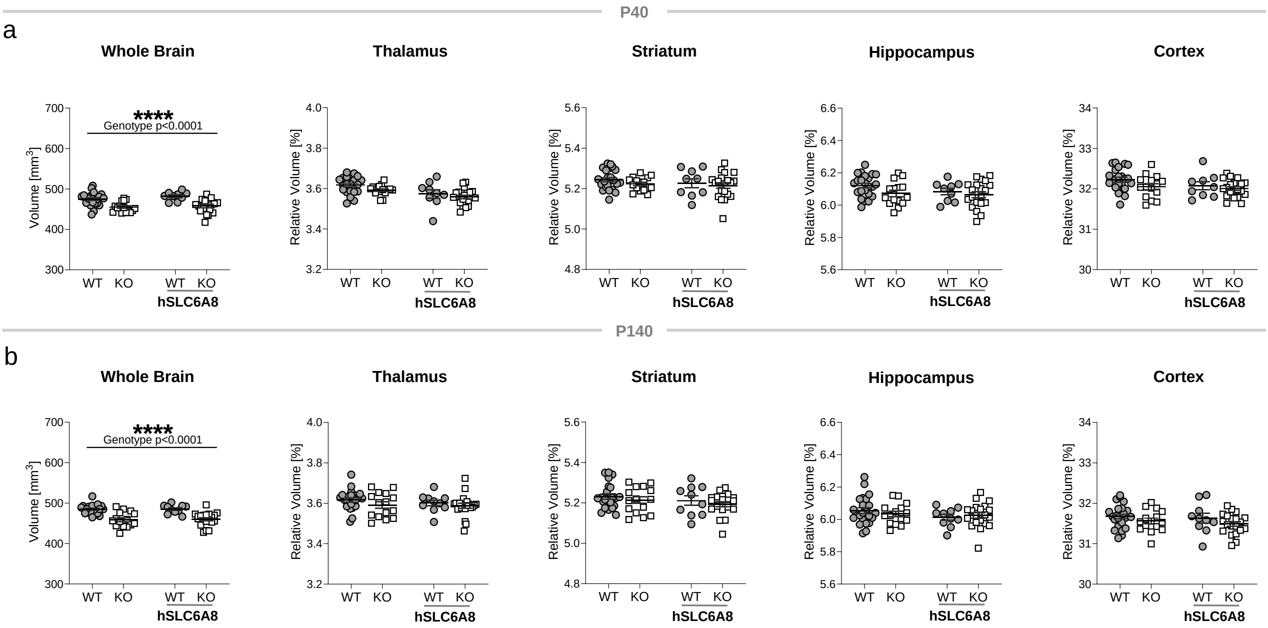


***Supplementary Figure 9. Reduced whole brain volume in Slc6a8-KO mice.*** *(**a**) Whole and regional brain volume quantification in Slc6a8-KO and WT mice at P40 (two-way ANOVA; genotype F = 30.96, p < 0.0001; genotype x treatment interaction: F = 0.32, p = 0.57). Region-specific volumes at P40 (Thalamus, two-way ANOVA; genotype F = 3.5, p = 0.066, interaction F = 0.17, p = 0.68; Striatum, genotype factor F = 1.41, p = 0.24, interaction F = 0.078, p = 0.78; Hippocampus, genotype factor F = 3.09, p = 0.08, interaction F = 0.59, p = 0.44; Cortex, two-way ANOVA, genotype factor F = 2.95, p = 0.09, interaction F = 0.51, p = 0.47). (**b**) Whole and regional brain volume quantification in Slc6a8-KO and WT mice at P140 (two-way ANOVA; genotype factor F = 44.5, p < 0.0001; interaction F = 0.0007, p = 0.98). Region-specific volumes at P140 (Thalamus, genotype factor F = 2.12, p = 0.15, interaction F = 0.18, p = 0.67; Striatum, genotype factor F = 1.18, p = 0.28, interaction F = 0.0009, p = 0.98; Hippocampus, genotype factor F = 0.04, p = 0.85, interaction F = 0.74, p = 0.39; Cortex, two-way ANOVA, genotype factor F = 2.89, p = 0.09, interaction F = 0.037, p = 0.85). Plots report absolute volume in* $mm^3$ *and relative volume % normalized to whole brain volume, error bars indicate SEM; dots represent individual values; ****p < 0.0001.*

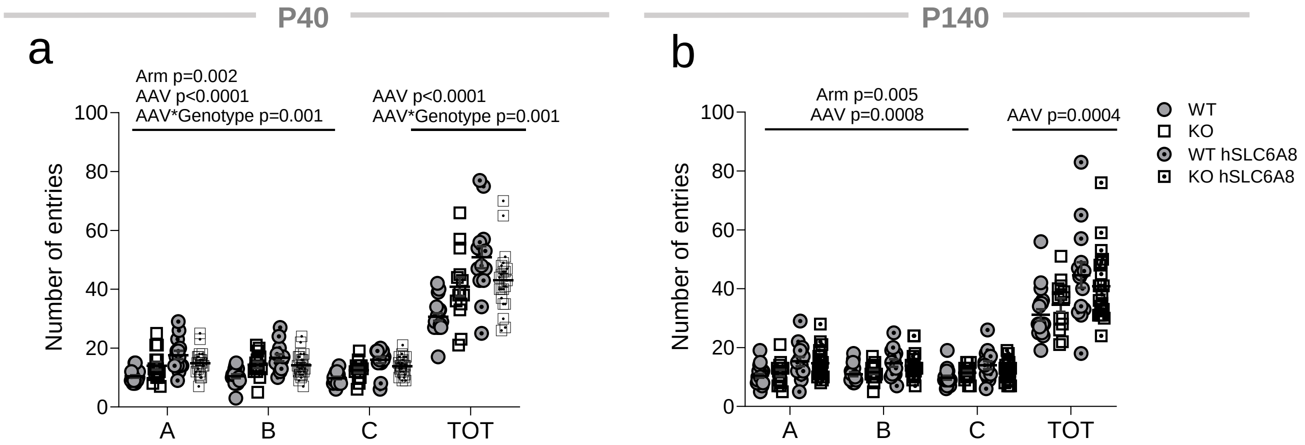


***Supplementary Figure 10. Behavioral performance following treatment with AAV-hSLC6A8.*** *Histograms depict the mean number of entries in the single arms of the maze (A, B, C) and the total number of arm entries (TOT) for the different experimental groups at P40 (**a**) and P140 (**b**). An increased number of entries was found in mice injected with AAV-hSLC6A8 both at P40 (single arms: Three-way RM ANOVA, factor AAV F = 18.8, p < 0.0001; TOT: two-way ANOVA factor AAV F= 20.05, p < 0.0001) and P140 (single arms: Three-way RM ANOVA, factor AAV F = 12.42, p = 0.0008; TOT: two-way ANOVA factor AAV F= 13.9, p = 0.0004).*

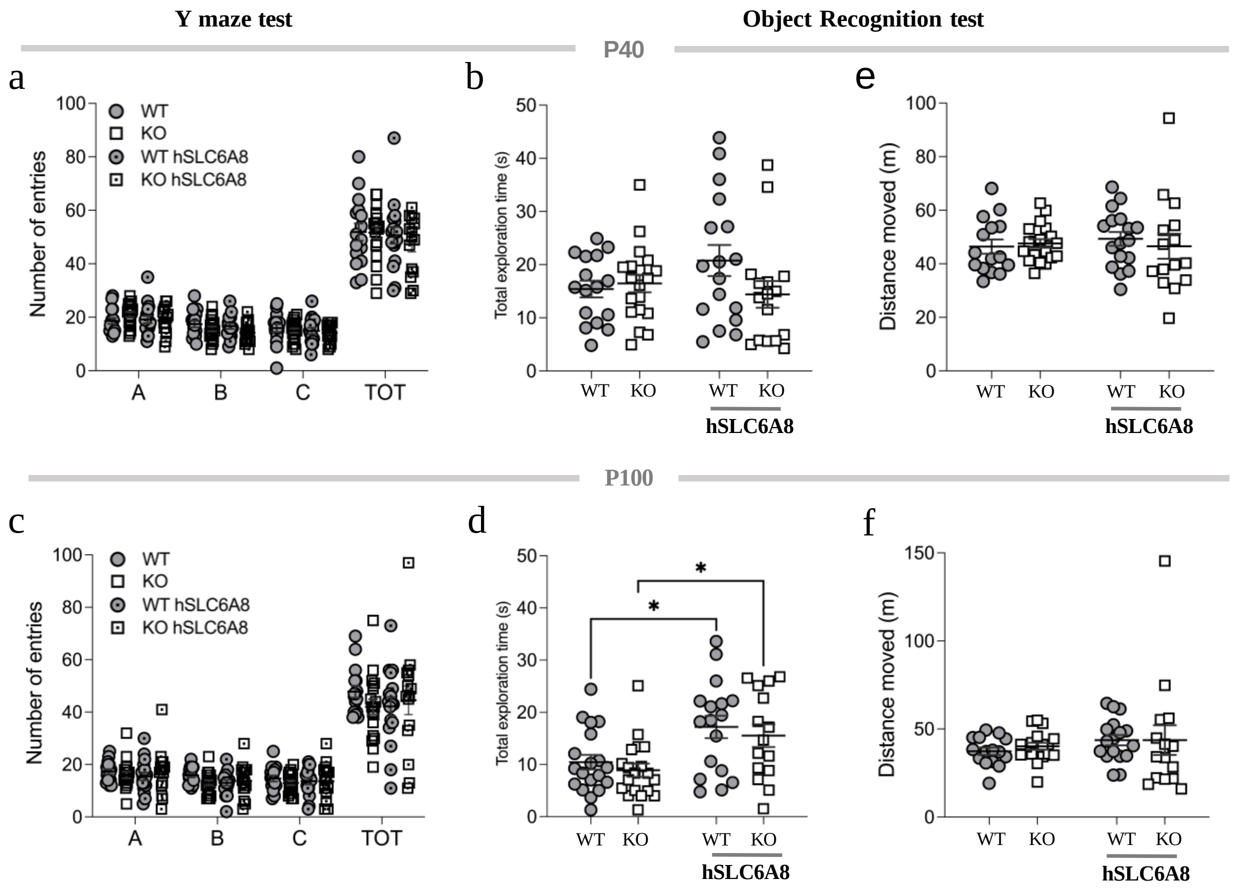


***Supplementary Figure 11. Perinatal AAV-hSLC6A8 injection does not alter overall exploratory behavior in WT and Slc6a8-KO mice.*** *(**a**) Number of entries in the Y-maze task (three-way ANOVA, interaction F = 0.540, p = 0.655) and (**b**) total exploration time in the Object Recognition Test (ORT) in the four experimental groups at P40 (two-way ANOVA, interaction F = 2.76, p = 0.102). At P100, the number of Y-maze entries remained unaffected across experimental groups (**c**, three-way ANOVA, interaction F = 1.15, p = 0.330), whereas object exploration time significantly increased in both genotypes following AAV-hSLC6A8 administration (**d**, two-way ANOVA, treatment F = 14.73, p = 0.0003). (**e-f**) Total distance moved in the ORT arena at P40 (e, two-way ANOVA, interaction F = 0.44, p = 0.51) and P100 (f, two- way ANOVA, interaction F = 0.101, p = 0.752). Error bars indicate SEM, each dot represents a mouse. Tukey's multiple comparison test, *p < 0.05.*

### Creatine Quantification

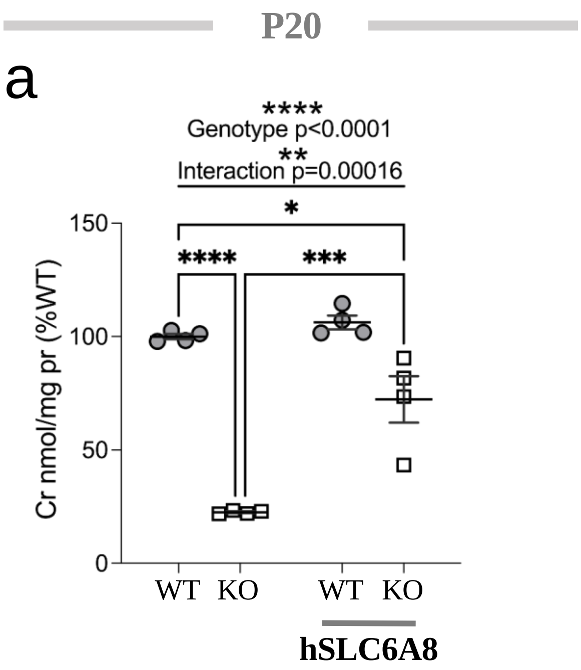


***Supplementary Figure 12. Quantification of brain creatine levels at P20*** *(two-way ANOVA, genotype factor $F = 107.1$ $p < 0.0001$, treatment $F = 27.1$, $p = 0.0002$, interaction $F = 16.3$, $p = 0.0016$). Error bars indicate SEM and each dot represents a mouse. Tukey's multiple comparison test, *$p < 0.05$, ***$p < 0.001$, ****$p < 0.0001$.*

**Table S1. Full sequence of pAAV-Jet-hSLC6A8**

cttccgcttcctcgctcactgactcgctgcgctcggtcgttcggctgcggcgagcggtatcagctcactcaaaggcggtaatacggttatccacagaatcaggggataacgcaggaaagaacatgtgagcaaaaggccagcaaaaggccagg
aaccgtaaaaaggccgcgttgctggcgtttttccataggctccgcccccctgacgagcatcacaaaaatcgacgctcaagtcagaggtggcgaaacccgacaggactataaagataccaggcgtttccccctggaagctccctcgtgcgctc
tcctgttccgaccctgccgcttaccggatacctgtccgcctttctcccttcgggaagcgtggcgctttctcatagctcacgctgtaggtatctcagttcggtgtaggtcgttcgctccaagctgggctgtgtgcacgaaccccccgttcagcccg
accgctgcgccttatccggtaactatcgtcttgagtccaacccggtaagacacgacttatcgccactggcagcagccactggtaacaggattagcagagcgaggtatgtaggcggtgctacagagttcttgaagtggtggcctaactacggcta
cactagaagaacagtatttggtatctgcgctctgctgaagccagttaccttcggaaaaagagttggtagctcttgatccggcaaacaaaccaccgctggtagcggtggtttttttgtttgcaagcagcagattacgcgcagaaaaaaaggatctc
aagaagatcctttgatcttttctacggggtctgacgctcagtggaacgaaaactcacgttaagggattttggtcatgagattatcaaaaaggatcttcacctagatccttttaaattaaaaatgaagttttaaatcaatctaaagtatatatgagtaa
acttggtctgacagttaccaatgcttaatcagtgaggcacctatctcagcgatctgtctatttcgttcatccatagttgcctgactccccgtcgtgtagataactacgatacgggagggcttaccatctggccccagtgctgcaatgataccgcg
agacccacgctcaccggctccagatttatcagcaataaaccagccagccggaagggccgagcgcagaagtggtcctgcaactttatccgcctccatccagtctattaattgttgccgggaagctagagtaagtagttcgccagttaatagtttg
cgcaacgttgttgccattgctacaggcatcgtggtgtcacgctcgtcgtttggtatggcttcattcagctccggttcccaacgatcaaggcgagttacatgatcccccatgttgtgcaaaaaagcggttagctccttcggtcctccgatcgttgtc
agaagtaagttggccgcagtgttatcactcatggttatggcagcactgcataattctcttactgtcatgccatccgtaagatgcttttctgtgactggtgagtactcaaccaagtcattctgagaatagtgtatgcggcgaccgagttgctcttgcc
cggcgtcaatacgggataataccgcgccacatagcagaactttaaaagtgctcatcattggaaaacgttcttcggggcgaaaactctcaaggatcttaccgctgttgagatccagttcgatgtaacccactcgtgcacccaactgatcttcagc
atcttttactttcaccagcgtttctgggtgagcaaaaacaggaaggcaaaatgccgcaaaaaagggaataagggcgacacggaaatgttgaatactcatactcttcctttttcaatattattgaagcatttatcagggttattgtctcatgagcgg
atacatatttgaatgtatttagaaaaataaacaaataggggttccgcgcacatttccccgaaaagtgccacctaaattgtaagcgttaatattttgttaaaattcgcgttaaatttttgttaaatcagctcattttttaaccaataggccgaaatcgg
caaaatcccttataaatcaaaagaatagaccgagatagggttgagtgttgttccagtttggaacaagagtccactattaaagaacgtggactccaacgtcaaagggcgaaaaaccgtctatcagggcgatggcccactacgtgaaccatcacc
ctaatcaagttttttggggtcgaggtgccgtaaagcactaaatcggaaccctaaagggagcccccgatttagagcttgacggggaaagccggcgaacgtggcgagaaaggaagggaagaaagcgaaaggagcgggcgctagggcgctggca
agtgtagcggtcacgctgcgcgtaaccaccacacccgccgcgcttaatgcgccgctacagggcgcgtcccattcgccattcaggctgcgcaactgttgggaagggcgatcggtgcgggcctcttcgctattacgccagctgcgcgctcgctc
gctcactgaggccgcccgggcaaagcccgggcgtcgggcgacctttggtcgcccggcctcagtgagcgagcgagcgcgcagagagggagtggccaactccatcactaggggttccttgtagttaatgattaacccgccatgctacttatcta
cgtagccatgctctaggaagagtaccattgacgtcaataatgacgtatgttcccatagtaacgccaatagggactttccattgacgtcaatgggtggagtatttacggtaaactgcccacttggcagtacatcaagtgtatcatatggggcggag
ttagggcggagccaatcagcgtgcgccgttccgaaagttgccttttatggctgggcggagaatgggcggtgaacgccgatgattatataaggacgcgccgggtgtggcacagctagttccgtcgcagccgggatttgggtcgcggttcttgtttg
tggggttcggcttctggcgtgtgaccggcggctctagagccaccatggcgaagaagagcgccgagaacggcatctatagcgtgtccggcgacgagaagaagggcccccctcatcgcgcccgggcccgacggggccccggccaagggcgacg
gccccgtgggcctggggacacccggcggccgcctggccgtgccgccgcgcgagacctggacgcgccagatggacttcatcatgtcgtgcgtgggcttcgccgtgggcttgggcaacgtgtggcgcttcccctacctgtgctacaagaacgg
cggaggtgtgttccttattccctacgtcctgatcgccctggttggaggaatccccattttcttcttagagatctcgctgggccagttcatgaaggccggcagcatcaatgtctggaacatctgtcccctgttcaaaggcctgggctacgcctccat
ggtgatcgtcttctactgcaacacctactacatcatggtgctggcctggggcttctattacctggtcaagtcctttaccaccacgctgccctgggccacatgtggccacaTctggaacactcccgactgcgtggagatcttccgccatgaagac
tgtgccaatgccagcctggccaacctcacctgtgaccagcttgctgaccgccggtcccctgtcatcgagttctgggagaacaaagtcttgaggctgtctggggggactggaggtgccaggggccctcaactgggaggtgaccctttgtctgctg
gcctgctgggtgctggtctacttctgtgtctggaagggggtcaaatccacgggaaagatcgtgtacttcactgctacattcccctacgtggtcctggtcgtgctgctggtgcgtggagtgctgctgcctggcgccctggatggcatcatttactat
ctcaagcctgactggtcaaagctggggtcccctcaggtgtggatagatgcggggacccagattttcttttcttacgccattggcctgggggccctcacagccctgggcagctacaaccgcttcaacaacaactgctacaaggacgccatcatc
ctggctctcatcaacagtgggaccagcttctttgctggcttcgtggtcttctccatcctgggcttcatggctgcagagcagggcgtgcacatctccaaggtggcagagtcagggccgggcctggccttcatcgcctacccgcgggctgtcacgc
tgatgccagtggccccactctgggctgccctgttcttcttcatgctgttgctgcttggtctcgacagccagtttgtaggtgtggagggcttcatcaccggcctcctcgacctcctcccggcctcctactacttccgtttccaaagggagatctctg
tggccctctgttgtgccctctgctttgtcatcgatctctccatggtgactgatggcgggatgtacgtcttccagctgtttgactactactcggccagcggcaccaccctgctctggcaggccttttgggagtgcgtggtggtggcctgggtgtacg
gagctgaccgcttcatggacgacattgcctgtatgatcgggtaccgaccttgcccctggatgaaatggtgctggtccttcttcaccccgctggtctgcatgggcatcttcatcttcaacgttgtgtactacgagccgctggtctacaacaacacc
tacgtgtacccgtggtggggtgaggccatgggctgggccttcgccctgtcctccatgctgtgcgtgccgctgcacctcctgggctgcctcctcagggccaagggcaccatggctgagcgctggcagcacctgacccagcccatctggggcct
ccaccacttggagtaccgagctcaggacgcagatgtcaggggcctgaccaccctgaccccagtgtccgagagcagcaaggtcgtcgtggtggagagtgtcatgtacccatacgatgttccagattacgcatgaggatccgagctcggtacca
agcttatcgataatcaacctctggattacaaaatttgtgaaagattgactggtattcttaactatgttgctccttttacgctatgtggatacgctgctttaatgcctttgtatcatgctattgcttcccgtatggctttcattttctcctccttgtataaa
tcctggttgctgtctctttatgaggagttgtggcccgttgtcaggcaacgtggcgtggtgtgcactgtgtttgctgacgcaacccccactggttggggcattgccaccacctgtcagctcctttccgggactttcgctttccccctccctattgcc
acggcggaactcatcgccgcctgccttgcccgctgctggacaggggctcggctgttgggcactgacaattccgtggtgttgtcggggaaatcatcgtcctttccttggctgctcgcctgtgttgccacctggattctgcgcgggacgtccttctg
ctacgtcccttcggccctcaatccagcggaccttccttcccgcggcctgctgccggctctgcggcctcttccgcgtcttcgccttcgccctcagacgagtcggatctccctttgggccgcctccccgcatcgataccgtcgacccgggcggcc
gcttcgagcagacatgataagatacattgatgagtttggacaaaccacaactagaatgcagtgaaaaaaatgctttatttgtgaaatttgtgatgctattgctttatttgtaaccattataagctgcaataaacaagttaacaacaacaattgcatt
cattttatgtttcaggttcagggggagatgtgggaggttttttaaagcaagtaaaacctctacaaatgtggtaaaatcgataaggatcttcctagagcatggctacgtagataagtagcatggcgggttaatcattaactacaaggaacccctagt
gatggagttggccactccctctctgcgcgctcgctcgctcactgaggccgggcgaccaaaggtcgcccgacgcccgggctttgcccgggcggcctcagtgagcgagcgagcgcgcagctgcattaatgaatcggccaacgcgcggggagag
gcggtttgcgtattgggcgct

**Table S2. Complete list of brain areas (and abbreviations) used for NBS.** 85 anatomically parcellated regions as in Coletta et al., 2020. ISO: isocortex, OLF: olfactory region, HPF: hippocampal formation, CTXsp: cortical subplate, STR:striatum, THAL: thalamus, HY: hypothalamus, MID: midbrain

| | | | | | | | |
|---|---|---|---|---|---|---|---|
| 1 | MOp | Primary motor area | ISO | 43 | PVR | Periventricular region | HY |
| 2 | MOs | Secondary motor area | ISO | 44 | MEZ | Hypothalamic medial zone | HY |
| 3 | Ssp | Primary somatosensory area | ISO | 45 | LZ | Hypothalamic lateral zone | HY |
| 4 | SSs | Supplemental somatosensory area | ISO | 46 | MBsen | Midbrain, sensory related | MID |
| 5 | GU | Gustatory areas | ISO | 47 | MBmot | Midbrain, motor related | MID |
| 6 | VISC | Visceral area | ISO | 48 | MBsta | Midbrain, behavioral state related | MID |
| 7 | AUD | Auditory area | ISO | 49 | MOp | Primary motor area | ISO |
| 8 | VIS | Visual area | ISO | 50 | MOs | Secondary motor area | ISO |
| 9 | ACAd | Anterior cingulate area, dorsal part | ISO | 51 | Ssp | Primary somatosensory area | ISO |
| 10 | ACAv | Anterior cingulate area, ventral part | ISO | 52 | SSs | Supplemental somatosensory area | ISO |
| 11 | PL | Prelimbic area | ISO | 53 | GU | Gustatory areas | ISO |
| 12 | ILA | Infralimbic area | ISO | 54 | VISC | Visceral area | ISO |
| 13 | ORB | Orbital area | ISO | 55 | AUD | Auditory area | ISO |
| 14 | AId | Agranular insular area, dorsal part | ISO | 56 | VIS | Visual area | ISO |
| 15 | AIp | Agranular insular area, posterior part | ISO | 57 | AId | Agranular insular area, dorsal part | ISO |
| 16 | AIv | Agranular insular area, ventral part | ISO | 58 | AIp | Agranular insular area, posterior part | ISO |
| 17 | RSPagl | Retrosplenial area, lateral agranular part | ISO | 59 | AIv | Agranular insular area, ventral part | ISO |
| 18 | RSPd | Retrosplenial area, dorsal part | ISO | 60 | PTLp | Posterior parietal association areas | ISO |
| 19 | RSPv | Retrosplenial area, ventral part | ISO | 61 | TEa | Temporal association areas | ISO |
| 20 | PTLp | Posterior parietal association areas | ISO | 62 | PERI | Perirhinal area | ISO |
| 21 | TEa | Temporal association areas | ISO | 63 | ECT | Ectorhinal area | ISO |
| 22 | PERI | Perirhinal area | ISO | 64 | PIR | Piriform area | OLF |
| 23 | ECT | Ectorhinal area | ISO | 65 | OLFnuc | olfactory_nuclei | OLF |
| 24 | PIR | Piriform area | OLF | 66 | CA | Ammon's horn | HPF |
| 25 | OLFnuc | olfactory_nuclei | OLF | 67 | DG | Dentate gyrus | HPF |
| 26 | CA | Ammon's horn | HPF | 68 | ENT | Entorhinal area | HPF |
| 27 | DG | Dentate gyrus | HPF | 69 | RHP | Retro_hippocampal | HPF |
| 28 | ENT | Entorhinal area | HPF | 70 | EndCla | EndopiriformNucleus_Claustrum | CTXsp |
| 29 | RHP | Retro_hippocampal | HPF | 71 | LA | Lateral amygdalar nucleus | CTXsp |
| 30 | EndCla | EndopiriformNucleus_Claustrum | CTXsp | 72 | BLA | Basolateral amygdalar nucleus | CTXsp |
| 31 | LA | Lateral amygdalar nucleus | CTXsp | 73 | BMA | Basomedial amygdalar nucleus | CTXsp |
| 32 | BLA | Basolateral amygdalar nucleus | CTXsp | 74 | PA | Posterior amygdalar nucleus | CTXsp |
| 33 | BMA | Basomedial amygdalar nucleus | CTXsp | 75 | STRd | Striatum dorsal region | STR |
| 34 | PA | Posterior amygdalar nucleus | CTXsp | 76 | STRv | Striatum ventral region | STR |
| 35 | STRd | Striatum dorsal region | STR | 77 | LSX | Lateral septal complex | STR |
| 36 | STRv | Striatum ventral region | STR | 78 | sAMY | Striatum-like amygdalar nuclei | STR |
| 37 | LSX | Lateral septal complex | STR | 79 | PAL | Pallidum | PAL |
| 38 | sAMY | Striatum-like amygdalar nuclei | STR | 80 | DORsm | Thalamus, sensory-motor cortex related | THAL |
| 39 | PAL | Pallidum | PAL | 81 | DORpm | Thalamus, polymodal association cortex related | THAL |
| 40 | DORsm | Thalamus, sensory-motor cortex related | THAL | 82 | LZ | Hypothalamic lateral zone | HY |
| 41 | DORpm | Thalamus, polymodal association cortex related | THAL | 83 | MBsen | Midbrain, sensory related | MID |
| 42 | PVZ | Periventricular zone | HY | 84 | MBmot | Midbrain, motor related | MID |
| | | | | 85 | MBsta | Midbrain, behavioral state related | MID |
