## Supplementary Methods for "Connectomic and behavioral alterations in creatine transporter deficiency are partially normalized by gene therapy"

**Author affiliations:**

[§] Contributed equally

[+] Contributed equally

### Materials and Methods

#### Ethical statement

Animal research was conducted in agreement with the Italian Law (DL 26/2014 of Italian Ministry of Health implementing EU 63/2010) and the recommendations in the *Guide for the Care and Use of Laboratory Animals* of the National Institute of Health. Animal projects were also reviewed and approved by the Animal Care Committee of the University of Trento, Istituto Italiano di Tecnologia (authorization no. 377/20, 07753.28), the Animal Care Committe of the Neuroscience Institute of Consiglio Nazionale delle Ricerche (authorization no. 507/2018-PR; 761/2024-PR) and the Italian Ministry of Health.

#### Animals

Slc6a8-KO mice (C57BL/6J background[1]) were generated by the European Molecular Biology Laboratory (EMBL). Wild-type littermates of the same background were used as controls. Given that CTD is an X-linked disorder, only male mice were used in this study. Slc6a8$^{x/-}$ females were crossed with wild-type male mice to generate Slc6a8$^{-/y}$ males (Slc6a8-KO, hereafter KO). Slc6a8$^{+/y}$ wild-type male littermates were used as controls (WT). Mice were maintained in a controlled environment with humidity set at 60 ± 10% and temperature maintained at 21 ± 1 °C. Food and water were readily accessible ad libitum. Animals were daily inspected to assess their health and well-being. Each standard cage (391 x 199 x 160 mm) provided nesting material and accommodated from 2 to 5 mice.

#### Molecular cloning

The full-length human sequence of *SLC6A8* (*hSLC6A8*) was cloned under the control of the small JeT promoter[2] into a pAAV_WPRE.SV40 using standard molecular biology techniques. An HA tag at the C-terminal of *SLC6A8* was used to facilitate the detection of the transgenic protein (pAAV_JeT-hSLC6A8-HA_WPRE.SV40). The JeT-hSCL6A8 sequence was synthesized by Twin Helix (Milano, Italy) as reported in Table S1. To verify that the HA tag does not interfere with transporter function, we also generated a version of the pAAV_JeT-hSLC6A8_WPRE.SV40 plasmid without the HA tag.

The human SLC6A8 coding sequence (isoform 1) was subcloned between the XbaI and BamHI restriction sites of the pUC57 vector (GenScript). Thus, briefly, both the pUC57-SLC6A8 insert and the pAAV_JeT-hSLC6A8_WPRE.SV40 backbone were digested with XbaI (NEB #R0145S) and BamHI (NEB #R0136S) at 37°C for 1 hour. Digestion products were resolved via electrophoresis on a 1% agarose gel, and bands corresponding to the correct fragment sizes were excised and purified using the QIAquick Gel Extraction Kit (QIAGEN #28706). DNA concentrations were subsequently quantified using a Nanodrop spectrophotometer. Ligation of insert and backbone was carried out overnight at 16°C using T4 DNA ligase (NEB #M0202S) at a 6:1 molar ratio (insert to backbone). The ligation reaction was used the next day to transform chemically competent E. coli DH5α cells by heat shock (30 min on ice, 45 s at 42°C, then 2 min on ice). Transformed cells were plated and cultured overnight at 37°C. Finally, plasmid DNA was isolated from selected bacterial colonies using the Wizard Plus SV Minipreps DNA Purification System (Promega #A1330).

### Cell culture and transfection

HEK293T cells were maintained in Dulbecco's Modified Eagle Medium (DMEM) supplemented with 10% fetal bovine serum (FBS) and 1% penicillin/streptomycin. Cell cultures were incubated at 37°C in a humidified atmosphere with 5% $CO_2$. For analysis of creatine (Cr) levels, cells were seeded at a density of $4 \times 10^5$ cells per well in 6-well plates. Upon reaching ~80% confluency, cells were transfected using Lipofectamine 2000 (ThermoScientific) at a 1:2 DNA/Lipofectamine ratio. Each well received 1.5 µg of plasmid DNA (JeT-hSLC6A8 or JeT-hSLC6A8-HA). The following day, cells were stimulated for 24 hours with 125 µM Cr dissolved in complete DMEM. After treatment, the cells were centrifuged at 400g for 5 minutes, and pellets were stored at -80°C before proceeding with the analysis of creatine levels.

### Viral preparation

Serotype 9 adeno-associated viral (AAV) vectors containing pAAV_JeT-hSLC6A8-HA_WPRE.SV40 were produced by the University of Pennsylvania Vector Core (Philadelphia, PA). Viral titration via genome copy number counting was performed using digital droplet PCR. As control, we used an AAV9 containing pAAV_JeT-GFP_WPRE.SV40.

### Experimental design

*In vivo* studies were carried out at Italian Institute of Technology (IIT, Rovereto, Italy) or at Neuroscience Institute of Consiglio Nazionale delle Ricerche (CNR, Pisa, Italy). Mice were randomly assigned to different experimental groups using a random number generator to minimize bias and ensure that each group represented a fair and unbiased sample of the population. All experimenters were blind to genotype and treatment groups. The sample size required for each experiment was estimated using a power analysis using G*Power software[3]. The effects size was estimated based on similar studies from the lab using analogous methods and treatment designs[4,5]. While this computation produced a target number of $n$ = 10 animal per group, we ended up using larger groups to match injection conditions across litters, and account for possible loss of animals during our longitudinal investigations owing to bias during behavioral habituation, or imaging artefacts.

Investigations at IIT (Gozzi laboratory) were carried out on four groups of male mice as follows: Slc6a8-KO (n = 20) and WT (n = 10) injected with AAV-hSLC6A8, and Slc6a8-KO ($n$ = 15) and WT ($n$ = 23) injected with AAV-GFP as controls. The four groups of mice were administered with AAV vectors via intracerebroventricular (i.c.v.) injection at P1. The four experimental groups were subjected to two sessions of fMRI scans, at P40 and P140. Forty-eight hours after each fMRI session mice underwent two behavioral tests. The testing order consisted of self-grooming scoring, followed by the Y maze test. At the end of the experimental schedule, a small group of Slc6a8-KO (*n* = 5) and WT (*n* = 2) mice were transcardially perfused with 4% paraformaldehyde and coronal brain sections were acquired with a confocal microscope to assess the expression pattern of the AAV-hSLC6A8 vector. The rest of the experimental mice were used for post-mortem Cr quantifications, which at the IIT site were carried out using Liquid Chromatography-Tandem Mass Spectrometry (LC-MS).

To corroborate and expand our behavioral investigations, we replicated the Y-maze experiments and conducted an Object Recognition Test (ORT) in a new group of mice treated with either AAV-hSLC6A8 or PBS at two distinct developmental time points (P40 and P100). These tests were conducted at the CNR site (Baroncelli lab, Pisa). Newborn mice (P1) were intraventricularly injected with either adeno-associated virus (AAV-hSLC6A8) or phosphate-buffered saline (PBS) as control. This study involved four groups of male mice: Slc6a8-KO (n = 15) and WT (n = 17) receiving AAV-hSLC6A8, and Slc6a8-KO (n = 18) and WT (n = 16) receiving PBS. Cognitive performance was evaluated at two time points, P40 and P100, using the Y-maze and the ORT to assess working and recognition memory, respectively. At P100, a subset of animals (n = 4 per group) underwent post-mortem creatine (Cr) quantifications, which at the CNR site were carried out using Gas Chromatography-Mass Spectrometry (GC-MS). GC-MS Cr measurements were also performed at P20 and P40 in separate groups of Slc6a8-KO (n = 4) and WT (n = 4) animals treated with AAV-hSLC6A8, along with corresponding control groups of Slc6a8-KO (n = 4) and WT (n = 4) animals.

Finally, an additional group of animals composed of Slc6a8-KO (n = 4) and WT (n = 4) mice treated with AAV-hSLC6A8, and Slc6a8-KO (n = 2), and WT (n = 2) mice receiving PBS as control, were sacrificed at P40 for analysis of transgene biodistribution and cellular targeting via stereological immunohistochemistry. These animals were bred and analyzed at the CNR site.

#### AAV injection in newborn mice

Intracerebroventricular injection of AAV was performed as previously described[6]. Briefly, at P1 pups were placed on ice for few minutes to induce hypothermic anesthesia and then moved onto a cooled stereotaxic frame. Coordinates were adjusted on the base of lambda point of *repere* (X, Y, Z) = (1, ±0.3, -2.0) mm. Pups received bilateral injections of 1 ul of AAV vector ($3x10^9$ vg/mouse) using a glass capillary connected by a tube to a Hamilton syringe placed on a micropump. The AAV solution was infused over a 60s period and the pipette was removed after a delay of 30s to prevent backflow. We allowed the pups to recover from the anesthesia under a heating bulb, before transferring them back to the mother cage.

#### Functional and anatomical magnetic resonance imaging (MRI)

Resting state fMRI (rsfMRI) data were acquired as extensively described in[7–10]. Briefly, mice were anesthetized with isoflurane (4% induction, 2% maintenance), intubated and artificially ventilated. During resting-state scans, isoflurane was replaced with halothane (0.7%) to obtain light sedation. To assess potential differences in the sensitivity of Slc6a8-KO animals to the anesthetic used, we measured the Minimal Alveolar Concentration (MAC), in an independent group of adult mice, n = 10 WT, n = 5 Slc6a8-KO, as previously described[11,12]. We used a 7T MRI scanner (Bruker), Bruker Paravision software version 6, a 72-mm birdcage transmit coil and a 4-channel solenoid coil for signal reception[4,11,13]. For each session, in vivo structural images were acquired with a fast spin-echo sequence (repetition time [TR] = 5500 ms, echo time [TE] = 60 ms, matrix 192 × 192, the field of view 2 × 2 cm, 24 coronal slices, slice thickness 500 µm). BOLD rsfMRI time series were acquired using an echo planar imaging (EPI) sequence with the following parameters: TR/TE = 1000/15 ms, flip angle 30°, matrix 100 × 100, field of view 2.3 × 2.3cm, 18 coronal slices, slice thickness 600 µm for 1920 volumes (total duration 32 minutes).

#### Analysis of fMRI timeseries

rsfMRI BOLD time series were preprocessed as previously described[4,10,11,14]. We removed the first 50 volumes of each timeseries to allow for signal equilibration. BOLD timeseries were then despiked, motion corrected and spatially registered to a common reference brain template. Mean ventricular signal (corresponding to the averaged BOLD signal within a reference ventricular mask) and motion traces of head realignment parameters (3 translations + 3 rotations) were regressed out from each time series. Finally, we applied a spatial smoothing (full width at half maximum of 0.6 mm) and a band-pass filter to a frequency window of 0.01-0.1 Hz.

Network based statistics (NBS)[10,15] was carried out by extracting fMRI signal in 85 anatomically parcellated regions (based on the Allen brain Atlas, Table S2). The list of reemployed regions can be found in[16]. We next computed an unpaired two-tailed Student's t test for each element of the corresponding correlation matrix separately ($t > 2.7$-$3.5$). FWER correction at the network level was performed using 5000 random permutations ($p < 0.05$) as implemented in the network-based statistics (NBS) package[10,15]. The chord plots show the 85 parcels clustered in 9 anatomical meta regions (Table S2). The size of each regional arc is proportional to the number of the parcels in the arc. The link width between two meta regions is proportional, relatively to the meta region size, to the ratio of edges connecting the two regions that significatively changed (blue, reduced number of connections; red, increased number of connections).

rsfMRI connectivity was also probed using seed-based analyses[4,17]. A seed region was selected to cover the areas of interest, based on NBS results (Supplementary Fig. 1). Voxel-wise intergroup differences in seed-based mapping were assessed using a 2-tailed Student's t test ($|t| > 2$, $p < 0.05$) and family-wise error (FWE) cluster-corrected using a cluster threshold of $p = 0.050$ as implemented in FSL (https://fsl.fmrib.ox.ac.uk/fsl/). The code used for preprocessing and analyzing mouse rsfMRI data is available at: https://github.com/functional-neuroimaging/rsfMRI-preprocessing; https://github.com/functional-neuroimaging/rsfMRI-global-local-connectivity; https://github.com/functional-neuroimaging/rsfMRI-seed-based-mapping

#### Brain volume analysis

To investigate whether CTD affects brain structure, we analyzed longitudinal anatomical T2-weighted images acquired in mice used for rsfMRI at P40 and P140. We aimed to measure both total brain volume and relative volume of anatomical regions of interests[18]. T2-weighted anatomical scans were manually skull-stripped using ITKSNAP. We used the nipype python library[19] wrapper for ants[20] registration to compute symmetric diffeomorphic transform[21] from dsurqe[22] reference template on individual images (sequentially estimating center of mass initialization, Rigid, affine, SyN). The transform was applied on four predetermined volumes of interest (Cortex, Hippocampus, Striatum, Thalamus) derived from the Paxinos/dsurqe ontology (mouse-brain-template repository[23]). Relative anatomical volumes were calculated by dividing the transformed volume of interest by the whole-brain volume of each individual mouse.

#### Immunohistochemistry

Animals were perfused transcardially with 4% paraformaldehyde (PFA) in phosphate buffer. Following perfusion, brains were post-fixed in PFA and subsequently cryoprotected in 30% sucrose in phosphate-buffered saline (PBS). Coronal brain sections (30 µm thick) were obtained using a freezing microtome and collected in PBS before immunohistochemical processing.

For biodistribution analysis, every 18$^{th}$ coronal section was systematically selected for immunohistochemical processing. These sections were incubated for 15 minutes with a quenching solution containing 50 mM NH4Cl in PBS, followed by three washes in PBS. Sections were subsequently incubated for 1 hour at room temperature in a blocking solution containing 5% Normal Goat Serum and 0.03% Triton-X, dissolved in PBS. After blocking, sections were incubated overnight at 4°C with a primary antibody against the HA tag (1:500, Abcam ab9110).

For double labeling experiments, sections were co-incubated overnight with primary antibodies targeting specific cell-type markers, including neuronal marker MAP2 (1:250, Merck M9942); astroglial marker GFAP (1:500, Synaptic Systems 173006), microglial marker Iba-1 (1:400, Wako 019–19,741), oligodendroglial marker BCAS1 (1:500, Synaptic Systems 445003) Antigen-antibody interactions were visualized using Alexa Fluor-conjugated secondary antibodies (1:400, Invitrogen). Following immunolabeling, sections were counterstained with Hoechst dye (1:500, Sigma), mounted onto microscope slides, and coverslipped using Vectashield mounting medium for fluorescence (Vector Laboratories Inc.).

#### Quantification of HA-Tag Biodistribution and analysis of co-localization

To quantify the biodistribution of HA-tag-positive pixels, imaging parameters—including laser intensity, gain, and offset—were optimized at the start of acquisition and maintained consistently throughout the imaging process. The right hemisphere of each section was imaged using a Zeiss laser-scanning Apotome microscope equipped with a 20× objective. Serial optical images were acquired at 1.38 µm intervals, generating 4 optical sections (covering a total depth of 6 µm) for each section. A total of 20 sections per animal were analyzed.

To enhance detection sensitivity, maximum intensity projections (MIPs) were generated from the four consecutive optical sections displaying the highest mean pixel intensity. These MIPs were then imported into ImageJ for further analysis. A threshold was established based on negative control HA-tag staining from PBS-injected animals. Subsequently, the area fraction of HA-positive pixels in the whole hemisphere was quantified. For more targeted analysis, regions of interest (ROIs) were drawn around specific areas (i.e., cortex, hippocampus, and striatum), and the area fraction of HA-positive pixels within these ROIs was measured. On average, 14 sections from the cerebral cortex, 10 from the striatum, and 6 from the hippocampus were analyzed per animal.

#### Behavioral tests

To reduce the potential circadian effects on behavioral performance, testing was carried out during the same time interval each day (1-5 pm). For the analysis of spontaneous self-grooming, mice were individually placed in an open field arena for 20 minutes (40 cm × 40 cm × 40 cm). Sessions were recorded and mice automatically tracked using the ANY-maze software. After a 10-min habituation period, the cumulative time spent by mice grooming themselves was scored for 10 min as an index of stereotypic behavior[24,25]. Spatial working memory was tested using the Y-maze, as previously described[1,25]. We utilized a Y-shaped maze featuring three symmetrical, grey, solid plastic arms positioned at 120-degree angles, each arm measuring 35cm in length, 5cm in width, and 15cm in height. Mice were placed at the center of the maze and given 8 minutes to freely explore. To prevent the accumulation of odors, the maze was cleaned with 10% ethanol between trials. All sessions were video-recorded and mice were automatically tracked using ANY-maze. An arm entry was defined as when all four limbs of the mouse were within the arm. A triad was constituted when a mouse made three consecutive entries into different arms of the maze. The number of arm entries and triads were recorded to compute the alternation percentage. This was achieved by dividing the number of triads by the number of possible alternations and then multiplying by 100.

We also assessed declarative memory using the object recognition test (ORT). The ORT apparatus consisted of a square arena (60 × 60 × 30 cm) made of poly(vinyl chloride) with black walls and a white floor. Animals were first acclimated to the arena and testing room during a single 10-minute habituation session The ORT procedure involved two phases:

a- Sample Phase: Two identical objects were placed in diagonally opposite corners of the arena (~6 cm from the walls). Mice were allowed to explore the objects freely for 10 minutes before being returned to their home cages. The objects were made of plastic, metal, or glass, and were heavy enough to prevent displacement by mice.

b- Testing Phase: Testing was conducted 24 hours after the sample phase. One of the previously encountered objects was replaced with a novel object, while the other was substituted by an identical copy of the original. Both objects were placed in the same positions as in the sample phase, and mice were allowed 5 minutes of free exploration.

Animal position was continuously recorded using a video tracking system (Noldus Ethovision XT). The total movement of the animal (Total exploration time) and the total distance covered

(Distance moved) were automatically computed. A discrimination index was computed as DI = (Tnew - Told)/(Tnew + Told), where Tnew is the time spent exploring the new object, and Told is the time spent exploring the old one[1].

#### Creatine measurements

*Liquid Chromatography-Tandem Mass Spectrometry (LC-MS, performed at IIT)*

Brain samples were homogenized in Phosphate Buffer Saline-Protease inhibitor (100:1) on ice. An aliquot of each brain homogenate was extracted (1:3) with cold $CH_3CN$ containing creatine-(methyl-*d3*) as internal standard and centrifuged at 21.100 x g for 20min at 4°C. A calibration curve was prepared in Phosphate Buffer Saline containing 20% $CH_3CN$. The calibrators were extracted as the brain homogenates. The supernatants of the extracted brain homogenates and calibrators were further diluted 100-fold with 2mM $NH_4OAc$ in H2O (pH 8), and analyzed by LC-MS/MS on a Waters ACQUITY UPLC-MS/MS system consisting of a triple quadrupole detector (TQD) mass spectrometer equipped with an electrospray ionization interface (ESI) and a photodiode array λ detector (PDA) from Waters Inc. (Milford, MA, USA). Electrospray ionization was applied in positive mode. Compound-dependent parameters as Multiple Reaction Monitoring (MRM) transitions and collision energy were developed for the parent compound and the internal standard. The analyses were run on an ACE Excel 2 C18 (150x2.1mmID) with an ACE Excel UHPLC Pre-column Filter at 40°C, using 2mM NH4OAc in H2O (pH 8) (A) and CH3CN (B) as mobile phase at 0.2mL/min. A linear gradient was applied starting at 0%B with an initial hold for 3.5min, then 0-80%B in 2min and 80-0%B in 0.1min, followed by a hold for 2.4min at 0%B. All samples were quantified by MRM peak area response factor in order to determine the levels of Cr in the brain samples. Data were expressed as ng Cr per mg brain (ng/mg brain) and normalized by WT Creatine content (%WT).

*Gas Chromatography-Mass Spectrometry (GC-MS, performed at CNR)*

HEK293T pellets and brain samples were homogenized in 0.7 mL of ice-cold PBS buffer (Sigma-Aldrich, Italy) using an ultrasonic disruptor (Microson Heat System, NY, USA). Samples were centrifuged at 600 × g for 10 min at 4°C. A 50 µl aliquot of the surnatant was taken for protein content assessment, while the remaining volume was used for Cr analysis, as previously described[25,26]. For Cr extraction, 200 µl of surnatant were mixed with 50 µl of saturated sodium hydrogen carbonate and 50 µl of a solution containing 2-phenylbutyric acid (internal standard, I.S.) in toluene (10 µg/ml, Merck, Italy). Subsequently, 1 ml of toluene and 50 µl of hexafluoro-2,4-pentanedione (Merck, Italy) were added to form bis-trifluoromethyl-pyrimidine derivatives. The reaction mixture was stirred overnight at 80°C. After incubation, the organic layer was centrifuged and dried under a steam of nitrogen. The residue was derivatized at room temperature using 100 µl of BSTFA+TMCS (Macherey-Nagel, Italy) prior to injection into the GC-MS spectrometer. GC analysis was conducted using an Agilent INTUVO 9000 GC system equipped with an HP5MS INTUVO capillary column (0.25 mm × 30 m, 0.25 µm film thickness) and an Agilent 5977C mass spectrometer (Agilent Technologies, Italy). The mass spectrometer was operated in electron ionization (EI) single ion monitoring (SIM) mode. The following m/z ions were used for metabolite quantification: 192 for the internal standard (I.S.); 258 for Cr. Data analysis was performed with MassHunter software. Data were expressed as nmol Cr per mg protein (nmol/mg pr) and normalized relative to the creatine content measured in wild-type (WT) samples (% WT).

#### Partial Least Square Correlation

We assessed brain-behavior relationship using partial-least square analysis, a multivariate data-driven statistical technique. This analysis aims to maximize the covariance between imaging readouts (in our case, individual seed-based connectivity maps of the anterior cingulate), behavioral scores (i.e. self-grooming and spontaneous alternations) and biochemical data (brain Cr levels) by identifying latent components (LC) that represent the optimal weighted linear combination of the original variables[27]. The null hypothesis that the observed brain-behavior relationship could be due to chance was verified via permutation testing (1,000 iterations) of behavioral and biochemical data matrix. To avoid LCs being trivially driven by inter-group differences, permutations were performed within groups as per previous guidelines[28]. The stability of the contribution of each brain and behavioral element was assessed via bootstrapping, where bootstrap resampling was performed within each experimental group to avoid patterns being driven by group differences[29]. We tested the existence of a significant correlation between behavioral indexes and fMRI connectivity in the cingulate fMRI network, a network system that is relevant for all the behaviors assessed[10]. Post-mortem brain Cr level assessed at P140 was modeled as a continuous variable in the behavioral matrix to probe whether the multidimensional relationship between connectivity and behavior could be affected by Cr levels.
